# Heterozygous deletion of SYNGAP enzymatic domains in rats causes selective learning, social and seizure phenotypes

**DOI:** 10.1101/2020.10.14.339192

**Authors:** D. Katsanevaki, SM. Till, I. Buller-Peralta, TC. Watson, MS. Nawaz, D. Arkell, S. Tiwari, V. Kapgal, S. Biswal, JAB. Smith, NJ. Anstey, L. Mizen, N. Perentos, MW. Jones, MA. Cousin, S. Chattarji, A. Gonzalez-Sulser, O. Hardt, ER. Wood, PC. Kind

## Abstract

Pathogenic variants in *SYNGAP1* are one of the most common genetic causes of nonsyndromic intellectual disability (ID) and are considered a risk for autism spectrum disorder (ASD). *SYNGAP1* encodes a <u>syn</u>aptic <u>G</u>TPase <u>a</u>ctivating <u>p</u>rotein that modulates the intrinsic GTPase activity of several small G-proteins and is implicated in regulating the composition of the postsynaptic density. By targeting the deletion of exons encoding the calcium/lipid binding (C2) and <u>G</u>TPase <u>a</u>ctivating <u>p</u>rotein (GAP) domains, we generated a novel rat model to study SYNGAP related pathophysiology. We find that rats heterozygous for the C2/GAP domain deletion (*Syngap^+/Δ-GAP^*) exhibit reduced exploration and fear extinction, altered social behaviour, and spontaneous seizures, while homozygous mutants die within days after birth. This new rat model reveals that the enzymatic domains of SYNGAP are essential for normal brain function and provide an important new model system in the study of both ID/ASD and epilepsy.

## Introduction

Pathogenic mutations in genes expressed early in development contribute significantly to neurodevelopmental disorders that manifest during childhood and persist through adulthood (Parikshak et al., 2013). Such disorders often result in global developmental delay, compromised cognition and other impaired behaviours including delayed motor function, delayed or absent language acquisition and communication, as well as limited adaptive skills. Large-scale exome sequencing studies indicate that *SYNGAP1* is one of the most prevalent recurring genes accounting for as many as 0.5-1% of individuals with neurodevelopmental disorders (Deciphering Developmental Disorders, 2015, 2017; Satterstrom et al., 2020). Individuals with *de novo* pathogenic mutations in *SYNGAP1* present with moderate-to-severe intellectual disability (ID) and autism spectrum disorder (ASD) (Hamdan et al., 2011; Hamdan et al., 2009). Mutations in *SYNGAP1* are also a risk factor for epileptic encephalopathies and almost all individuals with such mutations have co-occurring childhood epilepsy (Berryer et al., 2013; Carvill et al., 2013; Mignot et al., 2016; Parker et al., 2015; Vlaskamp et al., 2019; von Stulpnagel et al., 2015).

*SYNGAP1* encodes multiple isoforms of a multifunctional, synaptically enriched protein, SYNGAP, that is essential for development and survival (Chen et al., 1998; Kim et al., 2003; Kim et al., 1998; Knuesel et al., 2005; Komiyama et al., 2002). *Syngap* heterozygosity (*Syngap^+/-^*) in mice is associated with behavioural and neurological phenotypes including deficits in learning and memory, pronounced hyperactivity, as well as reduced threshold for induced seizures and spontaneous epileptiform activity (Berryer et al., 2016; Clement et al., 2013; Creson et al., 2019; Guo et al., 2009; Muhia et al., 2009; Nakajima et al., 2019; Ozkan et al., 2014; Sullivan et al., 2020).

SYNGAP isoform identity regulates its function and subcellular distribution (Araki et al., 2020; Gou et al., 2020; Li et al., 2001; McMahon et al., 2012). However, all isoforms share a central region comprised of a calcium/lipid binding domain (C2) and a <u>G</u>TPase <u>a</u>ctivating <u>p</u>rotein (GAP) domain that function together to regulate the intrinsic GTPase activity of the small G proteins Ras and Rap (Krapivinsky et al., 2004; Pena et al., 2008; Walkup et al., 2015). In addition to its GAP activity, SYNGAP also regulates synaptic strength and size through its role as a scaffolding molecule by restricting access to PSD95 PDZ domains (Walkup et al., 2016); its binding to PSD-95 also appears to regulate the phase transition of the postsynaptic density (PSD) (Zeng et al., 2016).

While SYNGAP has both enzymatic and scaffolding functions, it is not known how the alteration of these individual functions contribute to *SYNGAP1* related pathophysiology. Interestingly, although most pathogenic *SYNGAP1* variants identified to date result in premature termination or complete loss of protein, missense or in-frame mutations within exons encoding the C2 or GAP domain have been identified in at least 14 individuals with ID (Berryer et al., 2013; Deciphering Developmental Disorders, 2017; Mignot et al., 2016; Vlaskamp et al., 2019). This raises interesting questions about the extent to which the enzymatic function of SYNGAP is responsible for behavioural and physiological phenotypes associated with *SYNGAP1* haploinsufficiency. For example, is the C2/GAP domain necessary for survival? And do these domains regulate a wide-range of behavioural traits, indicating that loss of its enzymatic function is the main feature of *SYNGAP1* haploinsufficiency? Answers to these questions will be important for understanding mechanisms underlying clinical traits associated with pathogenic *SYNGAP1* variants and related rasopathies as well as for developing targeted treatments for these disorders. To test the role of the C2/GAP domains in behaviour and physiology independent of its scaffolding role, we generated a rat model in which *Syngap* C2 and GAP domains were deleted.

## Results

### A novel rat model of *SYNGAP1* haploinsufficiency

To address whether loss of the C2/GAP domain recapitulates traits associated with *SYNGAP1* haploinsufficiency, rats were generated with specific ablation of exons encoding these domains. To delete these regions selectively, zinc finger nucleases designed to target *Syngap* exons 8 to 12 (Figure **1A**) were microinjected into the pronucleus of fertilized, one-cell embryos, and then bred onto a Long-Evans (LE) background. A 3584bp selective deletion and 3bp insertion in one rat line were confirmed by sequencing, which resulted in a mutant protein that is 377 amino acids smaller than the original (Figure **1B**). Mutant protein expression was confirmed by immunoblotting of hippocampal homogenates and found to be located at synapses by immunoblotting of hippocampal synaptosome (SNS) fractions (Figure **1C** and **Supplementary Figure 1**); several mutant bands can be observed as would be predicted due to the presence of multiple SYNGAP isoforms (McMahon et al., 2012). Full-length SYNGAP protein levels in homogenates and SNS were reduced in heterozygous mutant (*Syngap^+/Δ-GAP^*) rats relative to wild-type (*Syngap^+/+^*; WT) (+/+_hom_: 1 ± 0.076; *+/Δ-GAP*_hom_: 0.415 ± 0.04; t_hom_(6)=6.846, *p*=0.0005; Figure **1D**; +/+_syn_: 1 ± 0.006; *+/Δ-GAP*_syn_: 0.5906 ± 0.082; t_syn(4)_=4.441, *p*=0.0113; Figure **1E**), while total SYNGAP (full length + mutant) was comparable between genotypes (**Supplementary Table 1**). While *Syngap^+/Δ-GAP^* rats appeared healthy, fertile and indistinguishable from WT littermates, homozygous rats (*Syngap^Δ-GAP/Δ-GAP^*) did not survive beyond P10 (Figure **1F**), suggesting that the C2/GAP domains are essential for postnatal viability.

**Figure 1.**
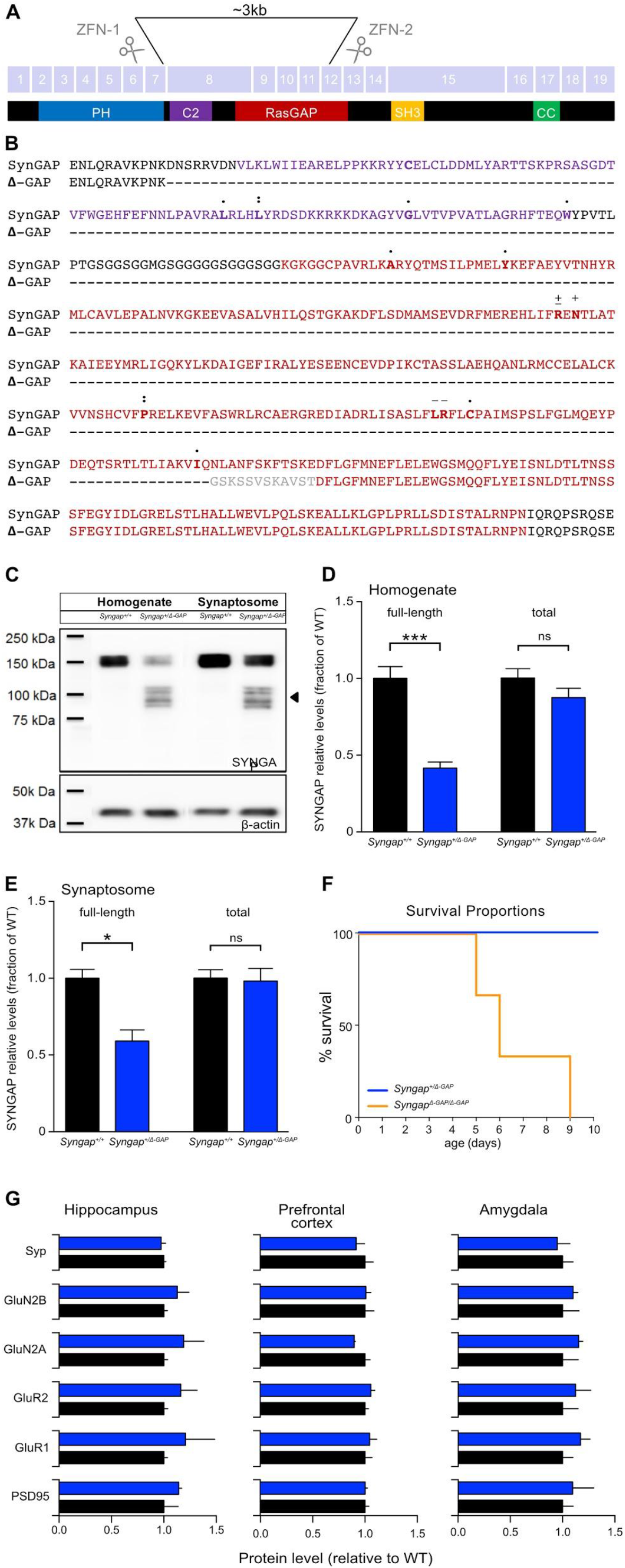
SYNGAP C2/GAP domain deletion strategy in rats results in reduction of endogenous SYNGAP expression and reduced viability. (A) Targeting strategy for ZFN-mediated selective deletion of *Syngap* exons 8-11 encoding C2/GAP domains. (B) Alignment of amino acids (aa) encoded by exons 8-13 of full-length and mutant SYNGAP proteins indicating the 377aa targeted deletion and unique aa resulting from 3bp insertion during targeting (grey) in relation to C2 (purple) and GAP (red) functional domains. + and - denote residues mutated to disrupt GAP function *in vitro* (Pena et al, 2008 and Vasquez et al, 2004, respectively); ± aa maps onto catalytic residue in p120GAP and NF1 (Ahmadian et al., 1997; Klose et al., 1998); dots indicate the number of instances and location of aa affected by missense mutations identified in these regions in individuals with MRD5. (C) Representative Western blot of extracts from rat hippocampal brain homogenates and synaptosomes. Bands in the molecular weight range expected for full length SYNGAP isoforms (~150kDa) were detected in homogenates and synaptosomes (SNS) from WT animals (Lanes 2 and 4, respectively). Additional bands (arrow) corresponding to the molecular weight range predicted for mutant SYNGAP isoforms are detected in homogenates and SNS from *Syngap^+/Δ-GAP^* rats. (D) Quantitation of full-length SYNGAP protein from homogenates reveals a significant decrease in *Syngap^+/Δ-GAP^* rats relative to WT while total SYNGAP (full length and mutant) is comparable to between genotypes; n_+/+_ = 4, n_*+/Δ-GAP*_ = 4. (E) As in homogenates, full-length SYNGAP protein is reduced in SNS from *Syngap^+/Δ-GAP^* rats but total protein levels are comparable to WT; n_+/+_ = 3, n_*+/Δ-GAP*_ = 3. mean ± SE is noted. (F) Juvenile *Syngap^Δ-GAP/Δ-GAP^* rats die by postnatal day 10 (n_*Δ-GAP/Δ-GAP*_ = 10). (G) Quantitation of pre- and post-synaptic proteins in SNS from hippocampus, prefrontal cortex and amygdala normalised to total protein and wild-type littermate controls.

Since SYNGAP plays a key role in synaptic modulation (McMahon et al., 2012; Walkup et al., 2016) we asked whether heterozygous C2/GAP domain deletion results in alterations in the molecular composition of synapses. Because SYNGAP is thought to regulate incorporation of glutamate receptors in the PSD (Rumbaugh et al., 2006; Vazquez et al., 2004; Walkup et al., 2016), we first compared the expression level of several proteins associated with postsynaptic function in purified SNS fractions from *Syngap^+/Δ-GAP^* rats and wild-type hippocampus, prefrontal cortex, and amygdala. Western blot analysis revealed no statistical differences in levels of post-synaptic proteins PSD95, AMPA receptor subunits GluA1 and GluA2, and NMDA receptor subunits GluN2A and GluN2B or of pre-synaptic protein synaptophysin (Syp) between genotypes in SNS from P60 animals (Figure **1G**, **Supplementary Figure 1** and **Supplementary Table 1**). This suggests that the SYNGAP scaffolding function is maintained in *Syngap^+/Δ-GAP^* rats.

### *Syngap^+/Δ-GAP^* rats display impaired extinction in a cued-fear conditioning paradigm

Pathogenic *SYNGAP1* mutations are associated with significantly limited cognitive ability and consequent deficits in adaptive functioning, with anecdotal caregiver reports of behavioural inflexibility. To investigate the effect of heterozygous loss of the SYNGAP C2/GAP domain on cognition and adaptive behaviour, we used a cued fear conditioning task (flashing light CS, footshock US) to examine acquisition, recall and extinction of fear memory (Figure **2**). Neither *Syngap^+/Δ-GAP^* nor WT littermates expressed freezing in the conditioning chamber before experiencing the first US. Both genotypes showed comparable freezing over 6 paired CS-US presentations during conditioning (2-way RM ANOVA, effect of CS presentation F_(5,105)_=54.87, *p*<0.0001; genotype F_(1,21)_=0.1912, *p*=0.6664; CS x genotype F_(5,105)_=1.368, *p*=0.2425; Figure **2A**).

**Figure 2.**
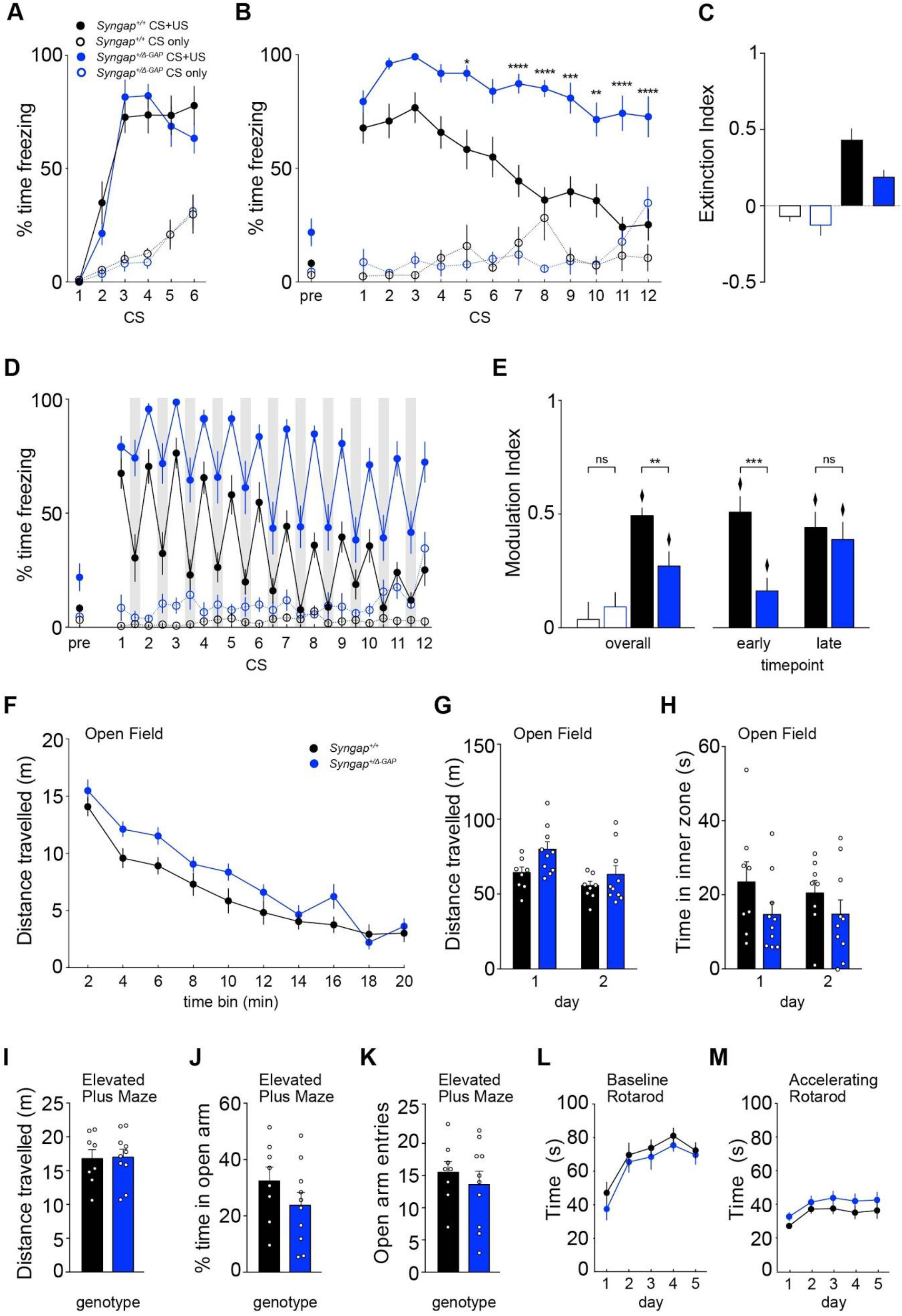
*Syngap^+/Δ-GAP^* rats display impaired extinction of fear association in a cued-fear conditioning paradigm. (A) During training, both WT and *Syngap^+/Δ-GAP^* rats display comparable levels of freezing to the flashing light that was paired with a mild foot shock (CS-US) (n_+/+ cs-us_ = 12, n_*+/Δ-GAP* cs-us_ = 11). (B) 24 hours after conditioning, *Syngap^+/Δ-GAP^* rats show increased fear responses to the neutral context but recall of fear memory to the first CS is comparable to WT. However, freezing to subsequent unreinforced CS presentations is significantly higher for *Syngap^+/Δ-GAP^* rats with the difference becoming more pronounced over consecutive presentations. In contrast, WT and *Syngap^+/Δ-GAP^* CS-only controls (n_+/+ cs-only_ = 7, n_*+/Δ-GAP* cs-only_ = 7) do not exhibit robust freezing to the CS during training (A) or recall testing (B). (C) Extinction index calculated as the change in % time freezing to the CS at the beginning and end of recall testing was significantly greater in conditioned WT than conditioned *Syngap^+/Δ-GAP^* rats or CS-only controls (n_+/+ cs-only_ = 7, n_*+/Δ-GAP* cs-only_ = 7, n_+/+ cs-us_ = 12, n_*+/Δ-GAP* cs-us_ = 11). (D, E) Comparison of the % time freezing *during* and *between* (shaded columns) CS presentations shows the CS specifically modulates freezing in conditioned WT and *Syngap^+/Δ-GAP^* rats (n_+/+ cs-only_ = 7, n_*+/Δ-GAP* cs-only_ = 7, n_+/+ cs-us_ = 12, n_*+/Δ-GAP* cs-us_ = 11). (F, G) *Syngap^+/Δ-GAP^* rats show an initial increase in locomotion during the first 20 min in the open field which habituates by day 2. (H) Time spent in the middle of the OF is comparable between genotypes (n_+/+ OF_ = 8, n_*+/Δ-GAP* OF_ = 10). (I-K) Behaviour of *Syngap^+/Δ-GAP^* rats was also indistinguishable from WTs during elevated plus maze testing, as indicated by locomotion, percentage of time spent in open arms (n_+/+ EPM_ = 8, n_*+/Δ-GAP* EPM_ = 10). (L, M) Motor coordination and learning is unaffected in *Syngap^+/Δ-GAP^* rats (n_+/+ RTR_ = 12, n_*+/Δ-GAP* RTR_ = 12). ITI: Inter-trial-Interval. *mean* ± SE is noted.

24 hours after training, rats were placed in a different testing context to assess retention and extinction of the conditioned response to unreinforced CS presentations. Both WT and *Syngap^+/Δ-GAP^* rats showed low freezing responses in the testing context prior to the first CS presentation, suggesting little fear generalization to the testing context (2-way ANOVA effect of genotype F_(1,33)_ = 3.667, *p*=0.0642; protocol F_(1,33)_ = 7.702, *p*=0.009; genotype x protocol F_(1,33)_ = 2.407, *p*=0.1303; Figure **2B**). Both groups showed similar high levels of freezing to the first presentation of the unreinforced CS during recall testing, suggesting comparable fear memory retention and expression. However, *Syngap^+/Δ-GAP^* rats showed more overall freezing during the recall test compared to WT rats which was due to greater freezing to subsequent unreinforced CS presentations (2-way RM ANOVA, effect of genotype F_(1,396)_ = 91.16, *p*<0.0001; also see **Supplementary Table 1**; Figure **2B**). Moreover, while WT rats decreased their freezing over repeated unreinforced CS presentations, *Syngap^+/Δ-GAP^* rats showed no apparent extinction learning (2-way ANOVA, effect of genotype F_(1,33)_ = 5.653, *p*=0.0234; CS F_(1,33)_ = 40.93, *p*<0.0001; genotype x CS F_(1,33)_ =2.198, *p*=0.1477; Figure **2C**), suggesting reduced behavioural flexibility in this associative learning task.

Control rats receiving unreinforced CS during training (CS-only) showed significantly less freezing than rats trained with the paired CS-US protocol (3-way ANOVA, effect of protocol F_(1,198)_ = 203.3, *p*<0.0001; CS x protocol F_(5,198)_ = 14.27, *p*<0.0001; Figure **2A**), implying that the flashing light of the CS is not aversive on its own. Rats trained with the paired CS-US protocol froze more during the CS presentations than between CS presentations regardless of genotype, whereas the CS did not phasically modulate freezing behaviour in control rats that received unreinforced CS during training (Figure **2D**); this is confirmed by calculation of a modulation index whereby a positive value indicates more freezing during the CS than in its absence (2-way ANOVA, effect of protocol F_(1,33)_ = 29.26, *p*<0.0001; Figure **2E**). Although *Syngap^+/Δ-GAP^* rats show less modulation of freezing by the CS overall (2-way ANOVA, effect of genotype x protocol F_(1,33)_ = 5.551, *p*=0.025), the modulation index of rats trained with the paired CS-US protocol was significantly greater than zero (one sample *t-test*, t_*WT*(11)_ =14.545, *p*<0.001; t_Δ-GAP(10)_ =4.340, *p*=0.001), indicating that greater freezing of *Syngap^+/Δ-GAP^* across the extinction trial was not due to generalised fear. Consistent with this, of rats trained with the paired CS-US protocol, *Syngap^+/Δ-GAP^* rats exhibited less modulation of freezing by the CS than WT early in the recall test, but modulation was comparable between genotypes later in the test (**Supplementary Table 1**).

Several reports indicate that mouse models of *SYNGAP1* haploinsufficiency exhibit hyperactivity and abnormal measures of anxiety (Berryer et al., 2016; Guo et al., 2009; Muhia et al., 2009; Nakajima et al., 2019; Ozkan et al., 2014) which can affect performance in, and confound the analysis of, tasks designed to study animal cognition (Crawley et al., 1997). Therefore, we determined whether deletion of the C2/GAP domain leads to anxiety, hyperactivity or locomotor abnormalities in our *Syngap^+/Δ-GAP^* rats, assessing their behaviour in the open field and elevated plus maze. Overall distance travelled in the open field was significantly greater in *Syngap^+/Δ-GAP^* rats compared to WT controls, but both groups showed a similar decrease in locomotion over the 20 min session (2-way RM ANOVA effect of genotype F_(1,16)_=5.660, *p*=0.0301; effect of time F_(9,144)_=60.04, *p*<0.0001; genotype x time F_(9,144)_=1.235, *p*<0.2782; Figure **2F**). Both groups also decreased their locomotor activity between the first and second day of exposure and distance travelled on day 2 was comparable between genotypes (2-way RM ANOVA, effect of day F_(1,16)_=16.34, p=0.0009; genotype F_(1,16)_=3.579, p=0.0768; interaction day x genotype F_(1,16)_=1.653, p=0.2169; Figure **2G**). These data suggest that, although *Syngap^+/Δ-GAP^* rats may be initially hyperactive in an open field, this rapidly normalises as they habituate to the environment. *Syngap^+/Δ-GAP^* and WT rats spent equivalent amounts of time in the centre of the open field, suggesting comparable anxiety levels in both groups (2-way RM ANOVA, effect of genotype F_(1,16)_=2.633, *p*=0.1242; day F_(1,16)_=0.1767, *p*=0.6798; interaction day x genotype F_(4,88)_=0.2019, *p*=0.6592; Figure **2H**). Similarly, spontaneous activity in the elevated plus maze as indicated by the distance travelled was comparable between *Syngap^+/Δ-GAP^* and WT littermates (unpaired *t-test*; t_(16)_=0.1149, *p*=0.9099; Figure **2I**). Both groups also presented with similar levels of anxiety as indicated by entries into open arms (unpaired *t-test*; t_(16)_=0.6892, *p*=0.5006; Figure **2K**) and the percentage of time spent in the open arms (unpaired *t-test*; t_(16)_=1.273, *p*=0.2212; Figure **2J**).

To test whether heterozygous C2/GAP deletion affects motor coordination or learning, we measured latency to fall from the rotating cylinder on both the fixed speed and accelerating versions of the rotarod test. Performance was indistinguishable between *Syngap^+/Δ-GAP^* and WT littermates (baseline Rotarod: 2-way RM ANOVA, effect of day F_(4,88)_=12.43, *p*<0.0001; genotype F_(1,22)_=1.606, *p*=0.2183; interaction day x genotype F_(4,88)_=0.1084, *p*=0.9793; accelerating Rotarod: 2-way RM ANOVA, effect of day F_(4,88)_=4.757, *p*=0.0016; genotype F_(1,22)_=2.528, *p*=0.1261; interaction day x genotype F_(4,88)_=0.0724, *p*=0.9903; Figure **2L, M**). Overall, these data indicate that both groups had similar balance, coordination and motor learning. Taken together, our findings from these different behavioural tasks suggest that heterozygous deletion of the C2/GAP domains does not affect basal levels of anxiety, activity, or motor coordination.

### *Syngap^+/Δ-GAP^* rats display normal spatial reference memory and reversal learning

To further investigate the effect of the loss of C2/GAP domain of SYNGAP on cognitive function and adaptive behaviour, we tested allocentric spatial learning in a hippocampus-dependent reference memory task in the water maze ((Morris et al., 1982); Figure **3A**). The task assesses the ability to use distal cues in order to navigate to a hidden escape platform in a circular pool. During acquisition, both WT and *Syngap^+/Δ-GAP^* rats showed similar learning of the hidden platform location over six days of training as indicated by the decrease in mean path length taken to reach the platform location (2-way RM ANOVA, effect of day F_(5,75)_=16.49, *p*<0.0001; genotype F_(1,15)_=2.845, *p*=0.1123; interaction day x genotype F_(5,75)_=1.849, *p*=0.1136; Figure **3B**). During the two probe trials (i.e. the first trials of days 3 and 6, respectively), the percentage of time spent in the platform zone increased for both genotypes (2-way RM ANOVA, effect of probe trial F_(1,15)_=7.246, *p*=0.0167; genotype F_(1,15)_=0.4692, *p*=0.5038; interaction probe trial x genotype F_(1,15)_=0.0002, *p*=0.9888; Figure **3C**), indicating that spatial learning and recall is intact in *Syngap^+/Δ-GAP^* rats.

**Figure 3.**
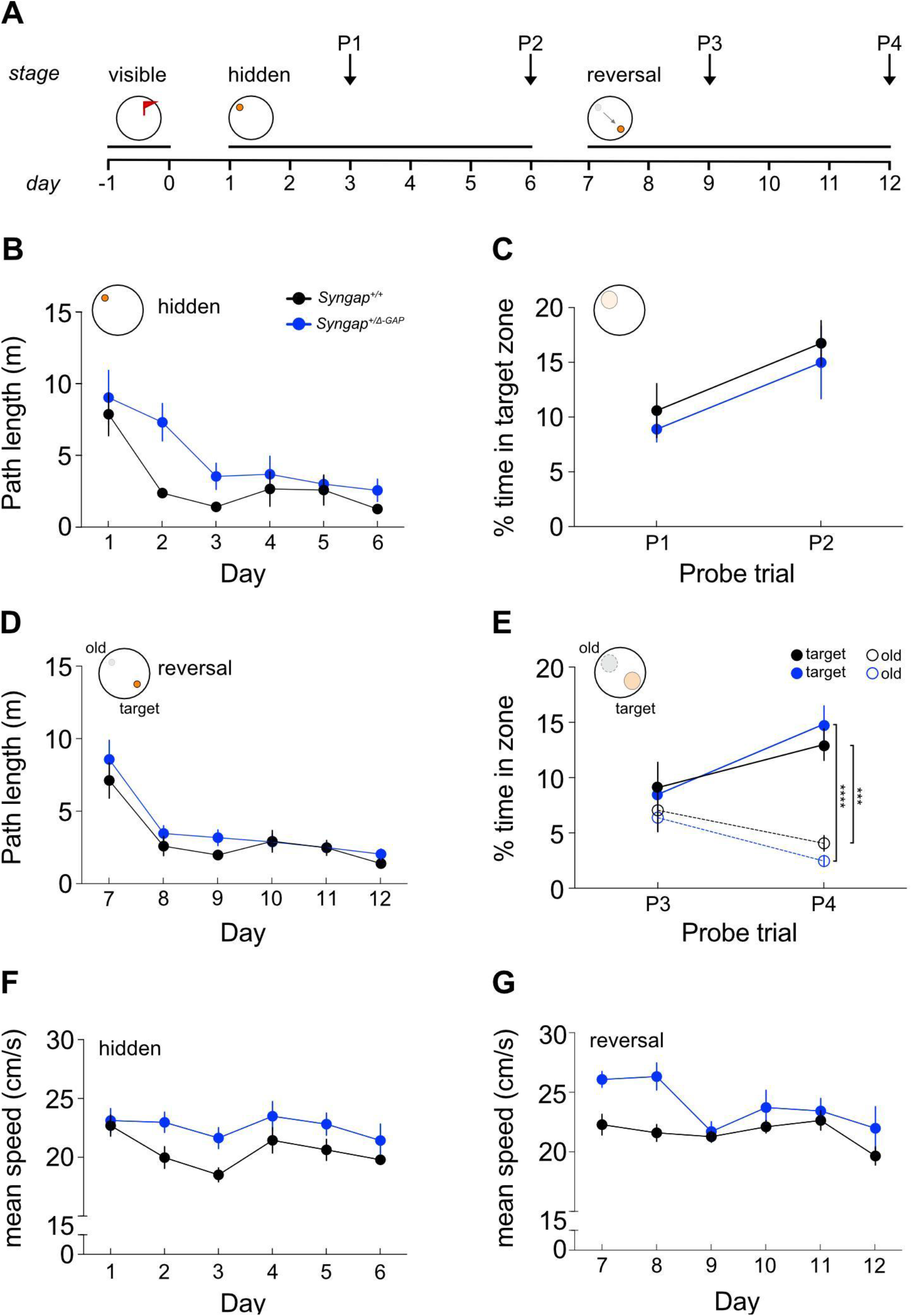
*Syngap^+/Δ-GAP^* rats have normal spatial reference memory acquisition and reversal learning. (A) Timeline of experimental protocol for spatial memory reference and reversal training in the water maze. (B) *Syngap^+/Δ-GAP^* rats learn the hidden-platform version of the water maze similarly to WTs as measured by a decrease over days in the path taken to escape and (C) the percent time in the platform location on daily probe trials. Performance during reversal learning was comparable between genotypes as measured by path to escape (D) and the percent time in the old and new platform locations during probe trials (E). (F, G) *Syngap^+/Δ-GAP^* rats swim faster than WT littermates overall (n_+/+_ = 8, n_*+/Δ-GAP*_ = 9). *mean* ± SE is noted.

We assessed reversal learning in this task as a measure of behavioural flexibility, by moving the platform to the opposite quadrant of the pool. Thus, in order to find the platform at the new location, animals need to stop visiting the old location of the platform while learning to swim to its new place. Reversal learning was equivalent between genotypes, as the path-length to reach the new escape location decreased similarly over days in both groups (2-way RM ANOVA, effect of day F_(5,75)_=29.48, *p*<0.0001; genotype F_(1,15)_=1.159, *p*=0.2987; interaction day x genotype F_(5,75)_=0.56, *p*=0.7303; Figure **3D**). Furthermore, in the reversal probe trials, both WT and *Syngap^+/Δ-GAP^* rats spent a higher percentage of time in the new target zone than in the previous platform location (‘old’) (Sidak’s multiple comparison test: for P4 *p_WT_*=0.0018, *p_HET_*<0.0001; see **Supplementary Table 1**; Figure **3E**), suggesting that behavioural flexibility was comparable between both groups in this spatial reversal task. While path length to escape was indistinguishable between genotypes during training and reversal learning, swim speed in the *Syngap^+/Δ-GAP^* rats was significantly higher than WT (2-way RM ANOVA, effect of genotype F_training(1,15)_=4. 945, *p*=0.0419; day F_training(5,75)_= 3.580, *p*=0.0059; interaction day x genotype F_training(5,75)_= 0.7567, *p*=0.5838; effect of genotype F_reversal(1,15)_=6.041, *p*=0.0266; day F_reversal(5,75)_=4. 714, *p*=0.0008; interaction day x genotype F_reversal(5,75)_= 1.885, *p*=0.1070; Figure **3F, G**). Similar to locomotion in the open field, this difference was only apparent at the start of training on each task (reference memory and reversal), so after day 3 of each task, the swim speed of *Syngap^+/Δ-GAP^* rats had decreased to the levels of WT rats. Together, these experiments indicate that *Syngap^+/Δ-GAP^* rats exhibit normal learning, recall, and behavioural flexibility in this spatial reference memory task.

### Altered social behaviour in *Syngap^+/Δ-GAP^* rats

Since impairments in social interactions are prevalent among ASD individuals, we used an adjusted three-chamber social interaction paradigm to assess social interaction and social preference ((Yang et al., 2011); Figure **4A**). After habituating the rats to the apparatus, we first assessed interaction with a caged, same-sex non-familiar WT conspecific compared to interaction with an empty cage. Sociability in this assay is typically defined as more time spent in the chamber with the non-familiar rat rather than in the other chamber, and more time spent sniffing the social than the non-social cage. Although *Syngap^+/Δ-GAP^* rats showed a decrease in overall exploratory behaviour (effect of genotype_chamber time_ F_(1,24)_=4.647, p=0.0414; Figure **4B**; and effect of genotype_sniffing time_ F_(1,24)_=24.55, p<0.0001; Figure **4D**), indices calculated using these measures indicate that both WT and *Syngap^+/Δ-GAP^* rats prefer to explore the social cage significantly more than would be expected by chance alone (see Supplementary Table 1; Figure **4C** and **4E**).

**Figure 4.**
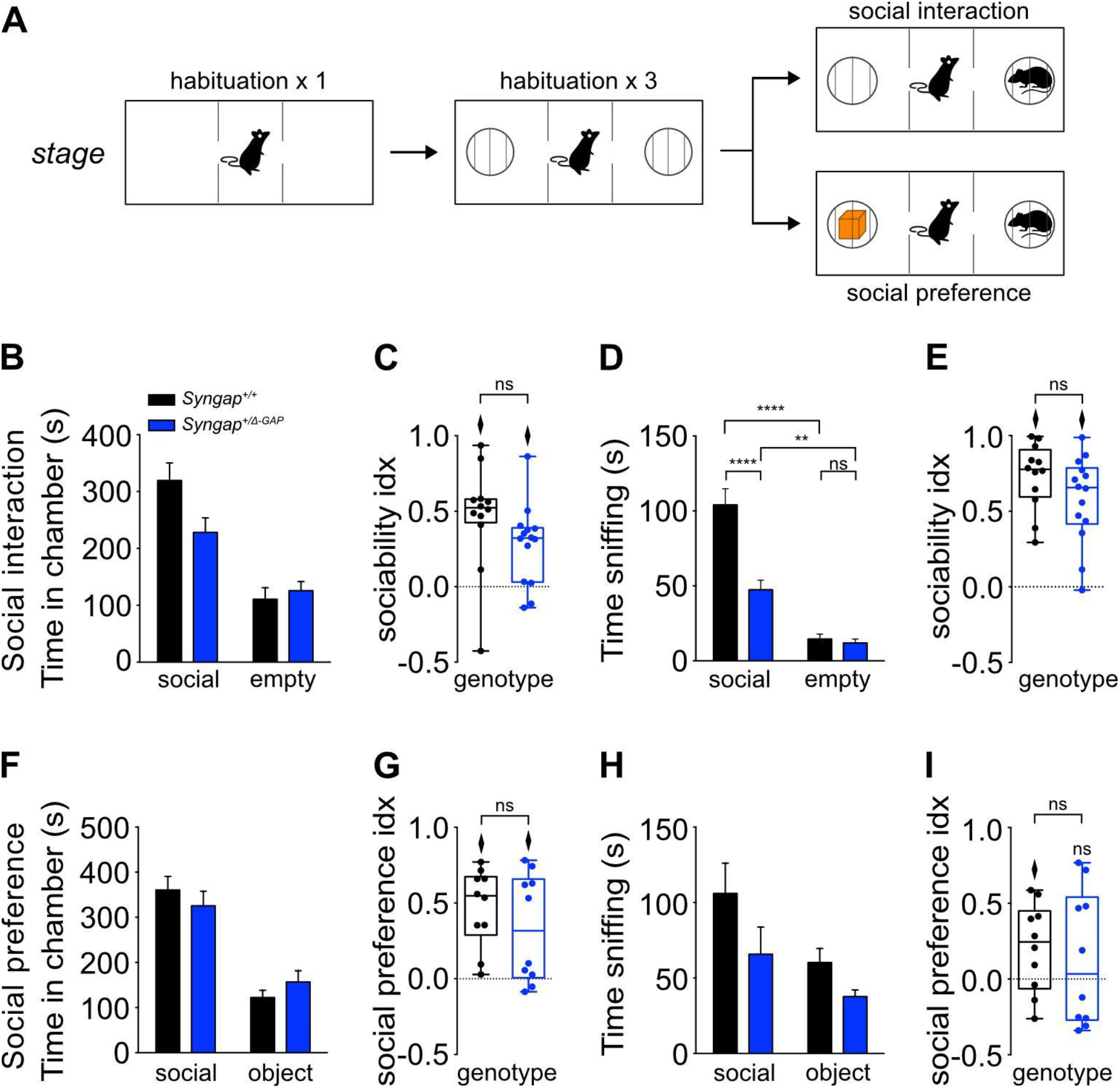
Altered social behaviour in *Syngap^+/Δ-GAP^* rats. (A) Schematic of the 3 chamber tasks. In the social interaction task, time in chamber (B) and sociability index (C) indicate WT and *Syngap^+/Δ-GAP^* rats show a preference for spending time in the chamber containing a caged social stimulus compared to the chamber containing an empty wire cage. (D) Time actively exploring (sniffing) and (E) sociability index for active exploration suggest that both WT and *Syngap^+/Δ-GAP^* littermates explore the social stimulus more (n_+/+_ = 12, n_*+/Δ-GAP*_ = 14). In the social preference task, time in chamber (F) and social preference index (G) indicate WT and *Syngap^+/Δ-GAP^* rats spend significantly more time in the chamber containing a caged social stimulus compared to the chamber containing a novel object. However, *Syngap^+/Δ-GAP^* rats do not show preference for actively exploring the social stimulus over the object (H-I) (n_+/+_ = 10, n_*+/Δ-GAP*_ = 10). Diamonds illustrate above chance performance (*p*<0.05). See also Supplementary Figure 3 for results during the novelty phase of the tasks, i.e the first 3 min.

As the empty cages were present during habituation and therefore rats were familiar with them prior to the test, we aimed to determine whether this preference was due to preference for a social stimulus per se or to a more general novelty preference. A separate cohort of rats was run in a modified configuration of the task to assess whether there was a preference for interacting with an unfamiliar rat over a novel inanimate object. In this task configuration, WT and *Syngap^+/Δ-GAP^* rats showed similar exploration of the social cage (time in chamber: genotype x stimulus F_(1,18)_=0.3826, *p*=0.5440, Figure **4F**; and time sniffing: genotype x stimulus F_(1,18)_=0.4159, *p*=0.5271; Figure **4H**). Both WT and *Syngap^+/Δ-GAP^* rats preferred to spend time in the chamber containing the social stimulus over the novel object, as indicated by a discrimination index significantly greater than zero (**Supplementary Table 1**; Figure **4G**). However, by the same measure, the *Syngap^+/Δ-GAP^* rats did not show a significant preference for actively exploring (sniffing) the social stimulus over the novel object (**Supplementary Table 1**; Figure **4I**). This was true for the entirety of the experiment (10 min) but also for the novelty phase, i.e the first 3 min (**Supplementary Figure 2A-H**). Together, these findings suggest that *Syngap^+/Δ-GAP^* rats lack preference for active interaction with social over non-social novel stimuli, but do prefer to be in the vicinity of social over non-social stimuli.

To determine whether an inability to detect object novelty prevents *Syngap^+/Δ-GAP^* rats from preferentially interacting with social versus non-social stimuli, we tested performance on a series of spontaneous recognition memory tasks (Kwon et al., 2006; Till et al., 2015) which assess the ability to discriminate novel from familiar objects (OR), changes in pairings of objects with context (OCR), object with place (OPR) and object with place and context (OPCR) over a short (2 min) retention interval (see schema in **Supplementary Figure 3A**). First, we assessed engagement with novel stimuli by examining the average time rats spend exploring objects during the first sample phase of each discrimination task. Compared to WT littermates, *Syngap^+/Δ-GAP^* rats tended to spend less time exploring novel objects, but this difference was not statistically significant (2-way RM ANOVA, effect of genotype F_(1,11)_=4.752, *p*=0.0519; task F_(3.587, 39.46)_=1.185, *p*=0.3309; interaction task x genotype F_(7,77)_=0.6130, *p*=0.7436; **Supplementary Figure 3B**). Consistent with this trend, *Syngap^+/Δ-GAP^* rats also show decreased exploration in another task, the marble interaction task, traditionally used to assess repetitive behaviours in rodent models of autism (Silverman et al., 2010). *Syngap^+/Δ-GAP^* rats display significantly decreased duration (unpaired *t-test*; t_(22)_=2.161, *p*=0.0419; **Supplementary Figure 3D**) and frequency of interaction with the marbles (unpaired *t-test*; t_(22)_=2.634, *p*=0.0152; **Supplementary Figure 3E**). Therefore, to eliminate the possibility that reduced exploration affected performance in the discrimination tasks, we imposed an object exploration criterion (see Methods) during the sampling/testing phase(s). When we only considered WT and *Syngap^+/Δ-GAP^* rats that had reached this criterion, their discimination index (which is a measure of preference to explore the novel stimulus configuration over the familiar configuration) of both groups was significantly greater than zero (which would reflect equal exploration of novel and familiar) in all four recognition memory tasks (i.e. OR, OCR, OPR, OPCR; **Supplementary Figure 3C, Supplementary Table 1**). Moreover, the discrimination index did not differ between genotypes for any task (**Supplementary Table 1**), suggesting that even complex associative recognition processes remain intact in SYNGAP mutant rats.

To control for the possibility that olfactory impairments prevent *Syngap^+/Δ-GAP^* rats from discriminating non-social and social odours, we tested both groups in a modified odour habituation-dishabituation task (Yang & Crawley, 2009). Although *Syngap^+/Δ-GAP^* rats explored most odours significantly less than their WT littermates on the first exposure (two-tailed unpaired t-tests; t_banana(19)_=5.568, *p*<0.0001; t_almond(18)_=5.213, *p*<0.0001; t_social1(12)_=2.427, *p*=0.0319; t_social2(11)_=0.5930, *p*=0.5652; **Supplementary Figure 4C and F**) both genotypes showed a progressive decrease in sniffing over repeated presentations of the same odour and increased sniffing levels when a novel odour was presented (**Supplementary Figure 4A, B, D, and E**). These data indicate normal function of the main olfactory system and vomeronasal organ and suggest that the altered social behaviour in *Syngap^+/Δ-GAP^* rats is not driven by an inability to discriminate among social and non-social odours.

### Behavioural and network analysis reveals the presence of seizure-like events in *Syngap^+/Δ GAP^* rats that can be suppressed by ETX

In addition to cognitive and behavioural symptoms, more than 85% of individuals with pathogenic *SYNGAP1* mutations exhibit epilepsy (Berryer et al., 2013; Hamdan et al., 2009; Klitten et al., 2011; Mignot et al., 2016; Pinto et al., 2010; Vlaskamp et al., 2019). Moreover, several patients with epileptic encephalopathy, a debilitating form of epilepsy with poor diagnosis due to refractory seizures and cognitive arrest, were found to carry *de novo* truncating mutations in *SYNGAP1* (Carvill et al., 2013). We noted home-cage behaviours associated with absence seizures among *Syngap^+/Δ-GAP^* rats, including head bobbing and occasional forelimb clonus and loss of balance (**Supplemental video 1**), which map directly onto low-level Racine stages used to rate seizure intensity (Racine, 1972). To verify that these behaviours represent seizures we recorded from chronically implanted 32-channel skull surface grid EEG probes (Figure **5A**) coupled with video and accelerometer recordings.

**Figure 5.**
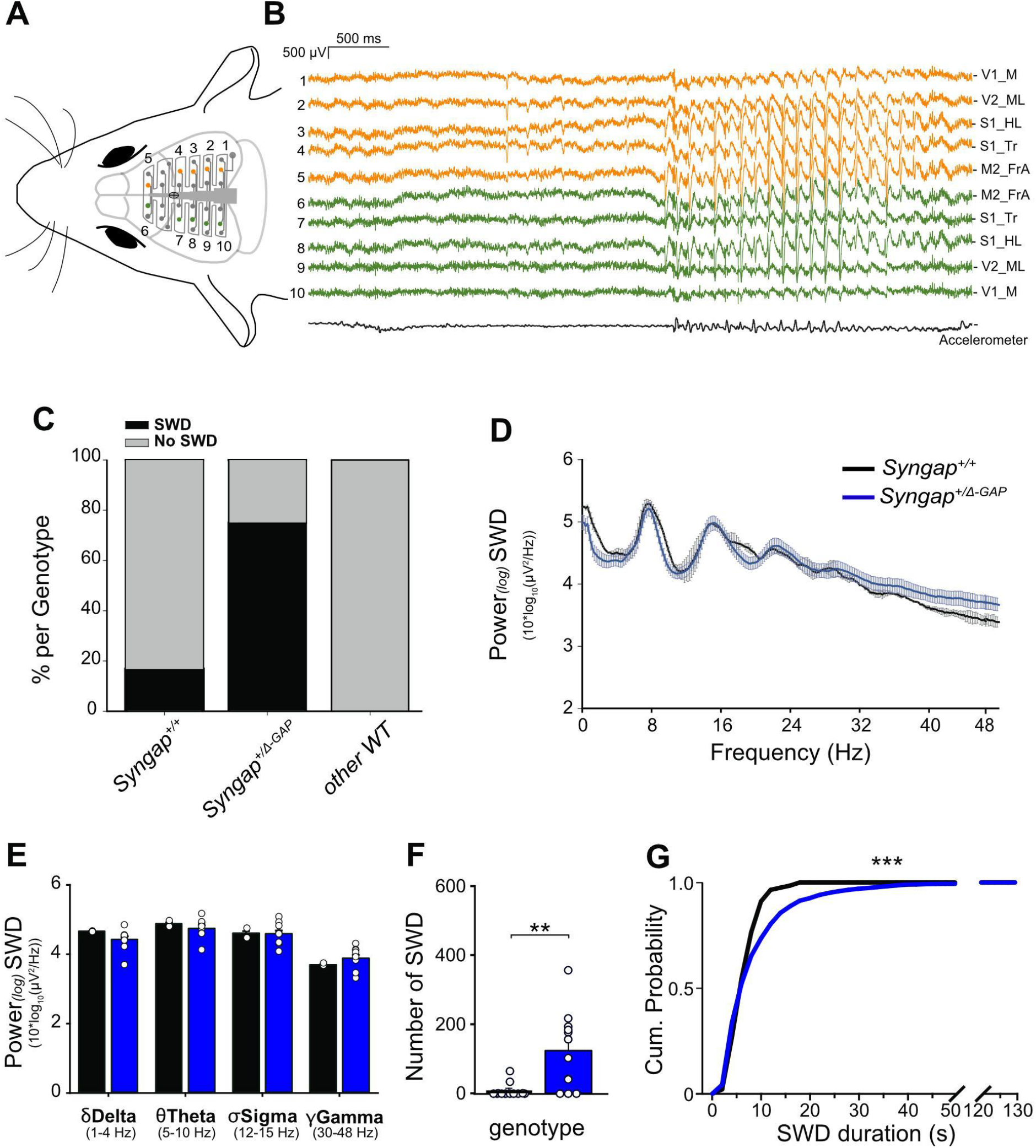
EEG analysis reveals the presence of SWDs in *Syngap^+/Δ-GAP^* rats. (A) Schematic of a 32-channel skull-surface EEG implant illustrating approximate location of electrodes relative to the brain and (B) representative traces from selected electrodes over both hemispheres (orange and green) reveal bilaterally occuring SWDs. (C) A significantly higher proportion of *Syngap^+/Δ-GAP^* rats exhibit SWDs than their WT littermates controls or WT rats from unrelated colonies (n_+/+_ = 12, n_*+/Δ-GAP*_ = 12, n_+/+other_ = 6). (D) SWD power from electrode overlaying S1 is comparable between *Syngap^+/Δ-GAP^* and WT littermates (n_+/+_ = 2, n_*+/Δ-GAP*_ = 9). (E) Averaged spectral power by EEG band for *Syngap^+/Δ-GAP^* and WT during SWDs (n_+/+_ = 2, n_*+/Δ-GAP*_ = 9). (F) The number of SWD events found in *Syngap^+/Δ-GAP^* and WT rats was significantly higher compared to WT littermates (n_+/+_ = 12, *n_+/Δ-GAP_* = 12). (G) Average cumulative probability of SWDs, by event duration in seconds, for *Syngap^+/Δ-GAP^* and WT rats suggests that SWD events detected in *Syngap^+/Δ-GAP^* rats were longer than those from WTs (from n_+/+_ = 2, n_*+/Δ-GAP*_ = 9). *mean* ± SE is noted. V1_M: primary visual cortex, medial component; V2_ML: secondary visual cortex, lateral component; S1_HL: primary somatosensory cortex, hindlimb region; S1_Tr: primary somatosensory cortex, trunk region; M2_FrA: secondary motor cortex, frontal association area.

Off-line visual and automated scoring identified prominent spike and wave discharges (SWDs) that generalized across all channels (Figure **5B**, **Supplementary Figure 5** for seizure detection). The incidence of SWDs in *Syngap^+/Δ-GAP^* rats was significantly higher (75%, 9/12 rats) than in WT littermates (16%, 2/12 rats), or in WT LE rats of the same age from an unrelated colony in the lab (0/6 rats) (Fisher’s exact test, *p*=0.014; Figure **5C**). In both *Syngap^+/Δ-GAP^* and WT littermates, SWDs co-occurred with an absence of locomotion and head bobbing related to absence seizures (**Supplemental video 2**). Spectral analysis of SWDs showed a prominent peak power in the theta band and a robust second harmonic (Figure **5D**) with no differences in power at any frequency bands between genotypes (Figure **5E**). The majority of SWDs occurred during quiet wakefulness (98.5% ± 0.6% *Syngap^+/Δ-GAP^* and 99.2% ± 0.8% WT) although the spectral properties of wakefulness were not significantly different between genotypes (**Supplementary Figure 5A-B**). Both the total number of SWDs (Figure **5F**) and number of SWDs per time awake were significantly higher in *Syngap^+/Δ-GAP^* rats compared to WT littermates (Mann-Whitney U test, *p*=0.002 and unpaired *t-test;* t_(22)_=3.794, *p*<0.001; **Supplementary Figure 5B and C** respectively). Moreover, cumulative frequency distribution profiles of SWD durations reveal that *Syngap^+/Δ-GAP^* rats had significantly longer SWDs than the WT littermates that exhibited SWDs (Kolmogorov-Smirnov test D(i30)=0.862, *p*<0.001; Figure **5G**).

Since SWDs are often associated with behavioural immobility in humans and rodents (Blumenfeld, 2005; Coenen & Van Luijtelaar, 2003), we tested whether the flashing lights used as a CS in the fear conditioning paradigm induced photosensitive SWDs that presented as enhanced behavioural immobility in *Syngap^+/Δ-GAP^* rats potentially confounding measures of freezing during fear recall and extinction (Figure **2A**). We recorded EEG from motor/parietal cortex and olfactory bulb while rats were introduced to flashing visual stimuli with the same properties as the CS previously used during the cued fear conditioning experiments (**Supplementary Figure 7A**). Flashing light exposure did not cause a change in the number of SWD events observed in either *Syngap^+/Δ-GAP^* rats or WT littermates (**Supplementary Figure 7B, C**) indicating that the observed increases in freezing during fear recall/extinction are not driven by CS induced seizures.

To assess whether the SWDs we observed are related to absence-like seizures, we evaluated whether they could be suppressed by ethosuximide (ETX), an T-type voltage-gated calcium channel antagonist commonly used to treat absence epilepsy in humans (Zimmerman & Burgemeister, 1958) and which blocks SWD in rodents (Terzioglu et al., 2006). EEG recordings over 5 consecutive days (Figure **6A**) revealed that a single dose of ETX significantly reduced SWD event number over a 2 hour period compared to no treatment or injection of saline alone (one-way RM ANOVA, effect of treatment F_(2,12)_=9.25, *p*=0.004; post-hoc paired t-tests - Holm-Sidak correction saline vs. ETX t_(6)_=4.25, *p*=0.003; untreated vs ETX t_(6)_=2.69, *p*=0.04; untreated vs saline t_(6)_=1.56, *p*=0.146; Figure **6B**). Seizure suppression by ETX was confirmed by calculation of a seizure index, whereby a negative value indicates fewer seizures than on the previous untreated day (one-way RM ANOVA, effect of treatment F_(3,18)_=18.24, *p*<0.001; post-hoc paired *t-test* - Holm-Sidak correction: ETX v pre-ETX vs post-ETX v pre-ETX *p*=0.004, ETX v pre-ETX vs sal v pre-sal *p*=0.006, ETX v pre-ETX vs post-sal v pre-sal *p*=0.005; Figure **6C**). The pharmacosensitivity of SWDs to ETX suggests the seizure-like events observed in *Syngap^+/Δ-GAP^* rats are related to absence epilepsy.

**Figure 6.**
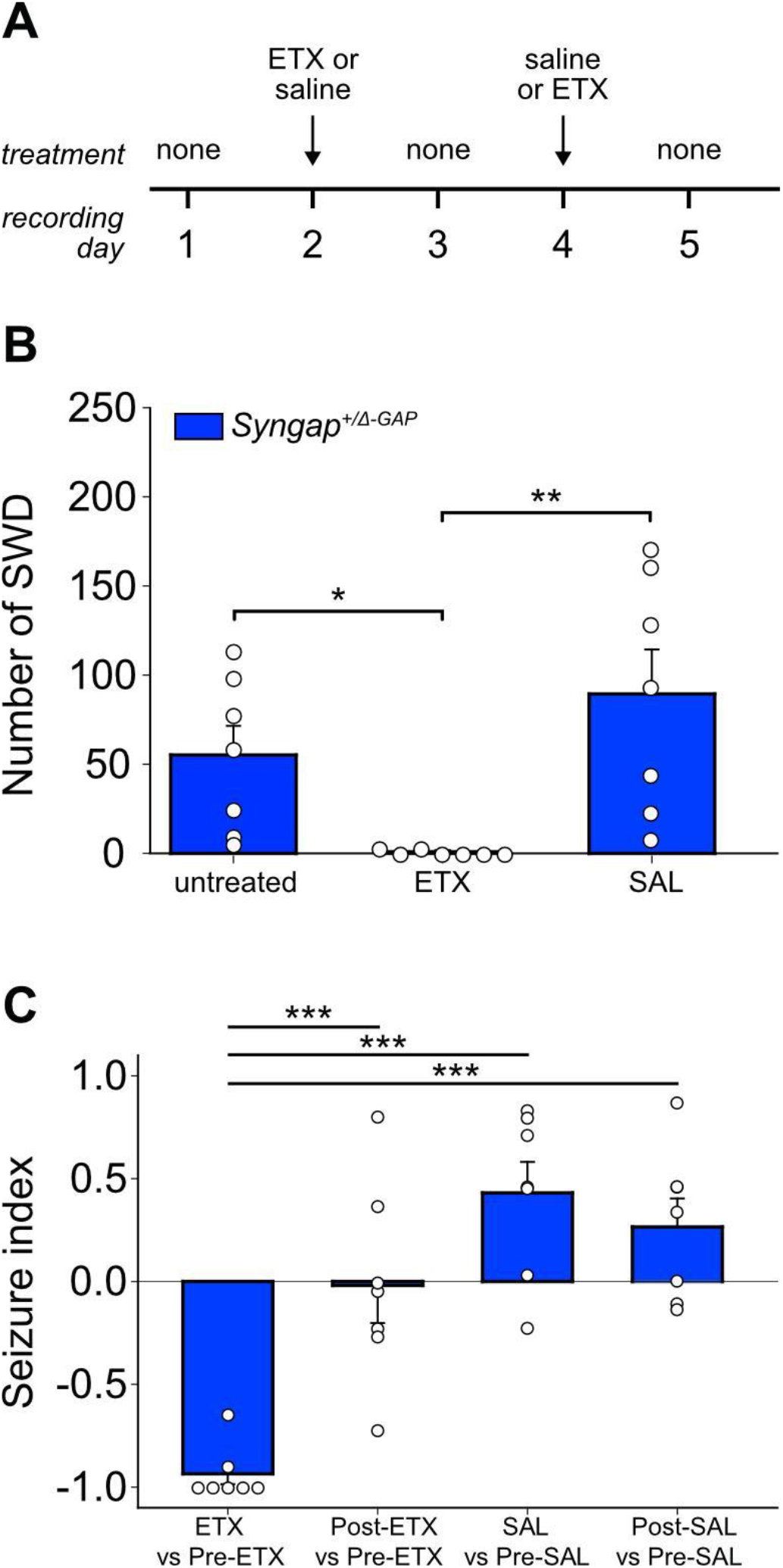
Ethosuximide reduces the number of SWDs in *Syngap^+/Δ-GAP^* rats. (A) Treatment timeline. (B) Number of SWD events identified in *Syngap^+/Δ-GAP^* rats after no treatment or following injection with ETX or saline (SAL) alone. (C) Seizure index compared to the previous untreated day shows greater suppression of SWD by ETX compared to other conditions (n_*+/Δ-GAP*_ = 7). *mean* ± SE is noted.

## Discussion

To test whether reduction in the GAP enzymatic activity of SYNGAP is key to clinical traits associated with *SYNGAP1* haploinsufficiency, we generated a rat model with a heterozygous deletion of the C2 and GAP domains of *Syngap*. Although overall total levels of SYNGAP expression were not affected, endogenous, full length SYNGAP was reduced to ~60% of WT levels in *Syngap^+/Δ-GAP^* rats. Importantly, the mutant protein localised to synapses and key post-synaptic proteins are present at normal levels in SNS from *Syngap^+/Δ-GAP^* rats enabling us to selectively test the role of the C2/GAP domains in behaviour and cognition independent of its scaffolding role. *Syngap^+/Δ-GAP^* rats demonstrate reduced exploration and fear extinction, altered social behaviour, and spontaneous seizures, indicating that many of the features of *SYNGAP1* haploinsufficiency result from a reduction in the regulation of the small G-proteins, Ras and Rap. Furthermore, the seizures and accompanying SWD are blocked by ETX, a drug commonly used to treat absence epilepsy suggesting a potential route to clinical benefit.

Using a range of behavioural tasks involving objects, social stimuli, and odours, we identified reduced exploration as a prevalent feature of *Syngap^+/Δ-GAP^* rats that is unlikely to have resulted from altered motor abilities or hyperactivity. Reduction in exploration is also unlikely to have resulted directly from absence seizures, since we did not observe behaviours characteristic of absence seizures during any of our behavioural tasks and seizures have previously been shown to be suppressed by mild sensory stimuli, for example those present during behavioural tasks (Pearce et al., 2014; Rodgers et al., 2015; Vergnes et al., 1982; Wiest & Nicolelis, 2003). Instead, the decrease in exploration may reflect an inability to maintain attention or a relative lack of interest and motivation to explore novel stimuli. Of note, mice with a genetic deletion of RICH2, a synaptic Rho-GAP that binds SHANK3, show a significant fear response to novel objects but above chance performance in object recognition (Sarowar et al., 2016). This suggests a more general involvement of modulators of small GTPases in behavioural responses to novelty.

Despite the decrease in exploration, we found that associative learning and fear learning were unaffected in *Syngap^+/Δ-GAP^* rats. However, they did exhibit a marked reduction in extinction of conditioned fear. This deficit in extinction learning did not result from a generalised increase in anxiety since we found no change in the open field or elevated plus maze. Furthermore, *Syngap^+/Δ-GAP^* rats exhibited modulation of their freezing to the CS. Together, this suggests heterozygous deletion of the GAP and C2 domains compromised the animals ability to learn that the conditioned stimulus no longer predicts the footshock. However, behavioural flexibility is not globally impaired in *Syngap^+/Δ-GAP^* rats since reversal learning in the watermaze was unaffected; this may indicate differential roles of SYNGAP across brain regions with circuits underlying emotional responses being particularly affected. Importantly, while discrimination of non-social objects and contexts was unaffected, we found that *Syngap^+/Δ-GAP^* rats display a significant impairment in exploring social over non-social stimuli. This may suggest that deletion of the C2/GAP domain of SYNGAP also results in other social impairments. Yet, we found that, like their WT littermates, *Syngap^+/Δ-GAP^* rats do prefer to interact with other rats rather than remaining alone. Because overall associative learning and memory also appears unaffected, it is possible that social preference impairments in *Syngap^+/Δ-GAP^* rats arise from altered sensory processing. In fact, recent studies in *Syngap^+/-^* mice highlighted dysfunction in the primary somatosensory cortex that is accompanied with abnormal tactile processing (Michaelson et al., 2018) and reports from clinicians indicate that sensory processing impairments are prevalent in individuals with pathogenic variants of *SYNGAP1* (Weldon et al., 2018). Alternatively, as both fear extinction and social behaviours involve the mPFC and amygdala in humans and rodents (Ko, 2017; Shin & Liberzon, 2010), alterations in these circuits could also contribute to these deficits. In support of this possibility, altered mPFC function has been described in *Syngap^+/-^* mice (Clement et al., 2013; Ozkan et al., 2014).

SWDs are a key electrophysiological feature in genetic rat models of absence seizures (Coenen & Van Luijtelaar, 2003; Shaw, 2004). SWDs were significantly more prevalent and had longer durations in *Syngap^+/Δ-GAP^* rats when compared to age-matched WT littermates, and similar to previous reports, these SWDs do not occur during our behavioural tasks and were suppressed by ethosuximide (Shaw, 2004, 2007; Terzioglu et al., 2006). Although SWDs were found in our WT animals, these were at a frequency similar to previous reports (Taylor et al., 2019). Moreover, the higher incidence of SWDs in mutants suggests that the deletion of the C2/GAP domain in rats drives cortical networks to a state of hyperexcitability leading to absence-like electrophysiological phenomena.

SYNGAP is a large, highly abundant synaptic protein with numerous isoforms arising from alternative promoter use and mRNA splicing (Chen et al., 1998; Kim et al., 1998; Li et al., 2001; McMahon et al., 2012). While a key role of SYNGAP is to regulate small G protein signalling, it also plays a key role as a scaffolding protein, anchoring AMPA receptors to the PSD through the regulation of transmembrane AMPA receptor-associated proteins (TARPs) and LRRTM2 (Walkup et al., 2016). Our finding that the Δ-GAP mutant protein, which maintains its PDZ binding domain, localises to synapses would appear to rule out the possibility that disruption to the scaffolding function contributes to the phenotypes observed in our animals. However, it should be noted that SYNGAP regulation of TARPs is a process that appears selectively in neurons from female rats (Mastro et al., 2020), while the majority of rats used in this study were males. Furthermore, SYNGAP has recently been demonstrated to form a homotrimer that binds PSD-95 to cause liquid phase separation of the PSD95-SYNGAP complex (Zeng et al., 2016). It has been proposed that this process mediates the association of SYNGAP with the PSD. Activity-dependent release from the PSD during stimuli that induce LTP cause the dispersal of SYNGAP, allowing AMPA receptor recruitment (Araki et al., 2015). However, whether this process happens *in vivo* or is important for the expression of clinical features of *SYNGAP1* haploinsufficiency is not known. SYNGAP dispersal from the PSD occurs even in the presence of RAS/RAP inhibitors (Araki et al., 2015) suggesting it is independent of the GAP domain function. Hence, the phenotypes presented here would be predicted to be independent of the role of SYNGAP in phase transition of the PSD. Ultimately, this would need to be directly tested with a mutation that prevents SYNGAP dispersal following LTP-induction while maintaining the ability of SYNGAP to regulate small G protein signalling.

Individuals with deleterious missense mutations in the C2 and GAP domains exhibit similar behavioural and neurological profiles to individuals with mutations predicted to lead to loss or truncation of the full length protein (Vlaskamp et al., 2019). This suggests that many of the clinical features of SYNGAP haploinsufficiency result from the decrease in the enzymatic function of SYNGAP. Our study supports an important role for the GAP and C2 domains, however a precise role for the enzymatic functions of SYNGAP in mediating behavioural and neurological phenotypes will require a direct comparison of the *Syngap^+/Δ-GAP^* rats with rats heterozygous for a null mutation in *Syngap*. While the behavioural domains that are affected in mice heterozygous for a null allele of *Syngap* and *Syngap^+/Δ-GAP^* rats are similar, the direction and quantitative nature of those changes appears to be quite different. However it is impossible to determine whether these reflect specific roles for the enzymatic domain or species specific differences in the expression of behaviours. Of note, *Syngap^+/-^* mice exhibit hyperactivity on a much greater scale (Berryer et al., 2016; Guo et al., 2009; Muhia et al., 2010; Nakajima et al., 2019; Ozkan et al., 2014) compared to that identified in *Syngap^+/Δ-GAP^* rats. Hyperactivity is a potential confounding factor in measuring performance in tasks designed to study animal cognition, including expression of defensive behaviours used in fear conditioning and other behavioural and cognitive phenotypes reported in *Syngap^+/-^* mice. What is clear is that the enzymatic domain is essential for survival since homozygous deletion of the C2/GAP domains results in perinatal lethality, similar to *Syngap* homozygous null mice (Kim et al., 2003; Knuesel et al., 2005; Komiyama et al., 2002) and rats (Mastro et al., 2020).

*SYNGAP1* haploinsufficiency is a complex disorder and further research will be necessary to identify how the other functions of SYNGAP may contribute to human pathophysiology. Our findings from a new rat model provide valuable insight into the phenotypic spectrum associated with mutations in the *SYNGAP1* gene in human patients of ID and further reinforces the need for more animal models in the field of neurodevelopmental disorders. Using a novel rat model, we demonstrate that disruption of the enzymatic domain of SYNGAP is a major contributor to the pathophysiology associated with *SYNGAP1* haploinsufficiency, providing key insight into potential therapeutic strategies. Further studies into the pathology associated with mutations that affect the scaffolding functions of SYNGAP will be required to further dissect the contribution of the structural properties of SYNGAP to the varied features of this disorder.

## Methods

### Animals

Subjects were Long Evans-*SG^em2/PWC^*, hereafter referred to as *Syngap^+/Δ-GAP^* bred in-house and kept in a 12h/12h light dark cycle with ad libitum access to water and food. Colony founders were produced by Sigma Advanced Genetic Engineering (SAGE) Labs (St. Louis, MO, US) using zinc finger nuclease (ZFN)–mediated deletion (Geurts et al., 2009) of the GAP domain of *Syngap*. Pups were weaned from their dams at postnatal-day 22 (P22) and housed in mixed genotype cages with littermates, 2-4 animals per cage. Animals were genotyped by PCR. 3-6 month old male/female animals were subsequently used for all experiments.

### RNA isolation and RT-PCR

Total hippocampus RNA was isolated from 4-month olds rats using RNeasy Lipid Tissue Kit (Qiagen) as per manufacturer’s instructions. 2 μg total RNA was used for cDNA synthesis using SuperScriptIII (Invitrogen) with oligo(dT) and random hexamers. PCR was performed using GC-RICH PCR System (Roche). *SYNGAP* primers (rSG_F: ATG ACC GGG CCC GGC TG and rSG_R: CTT CAG GAG GGC TTC CTT GCT GAG CT) spanning exons 5/6 and 12/13 boundaries, respectively with endogenous amplicon ~1630bp and mutant amplicon ~363 bp. Samples were run on a 1.0% agarose gel and gel purified prior to Sanger sequencing. Protein sequences were aligned using ClustalW; location of functional domains predicted using SMART (Letunic & Bork, 2018).

### Tissue preparation and immunoblotting

Hippocampi were dissected in ice cold ACSF from P60 *Syngap^+/Δ-GAP^* and WT littermates, snap frozen and stored at −80C until SNS preparation. Total tissue lysate was prepared in ice cold 1XSucrose/EDTA buffer (0.32M Sucrose, 1mM EDTA, 5mM Tris, PH 7.4) using 5-6 up-and-down strokes of a pre-chilled motorized Teflon glass homogenizer, followed by centrifugation at 1075 *g* for 10 minutes at 4°C. Pure synaptosomes (SNS; pinched off nerve terminals) were prepared by layering supernatant gently on top of a discontinuous Percoll-density gradient (3% uppermost, 10% middle, and 23% bottom; Percoll, P1644, Sigma-Aldrich, UK) and centrifuged at 47,807 *g* for 8 min at 4°C. The fraction between 23% and 10% was collected and re-suspended in HEPES-Buffered-Krebs (HBK-118.5mM NaCl, 4.7mM KCl, 1.18mM MgSO_4_, 10mM Glucose, 1mM Na_2_HPO_4_, 20mM HEPES, PH 7.4 balanced with Trizma) and SNS were pelleted out by centrifugation at 20,198 *g* for 15 min at 4°C. Homogenates were prepared from total tissue lysate by centrifugation at 25,128 *g* for 30 min. SNS pellets and homogenates were dissolved in RIPA buffer containing protease inhibitors (Roche complete mini EDTA-free protease inhibitor cocktail 4693159001, Sigma-Aldrich, UK) and phosphatase inhibitors (cocktail II P5726, Cocktail III P0044, Sigma-Aldrich, UK); proteins were estimated by MicroBCA Assay (Pierce BCA protein estimation kit, 23225, Thermofisher, UK).

Approximately 10μg of each protein extract was separated on a precast gradient gel (NuPAGE 4-12% Bis-Tris Protein Gels, NP0336BOX, Thermofisher) and transferred to PVDF membrane (GE10600022, Thermofisher, UK). The membrane was then blocked with 5% milk (Blotting grade blocker, 1706404, Bio-Rad) in TBST 1X at RT for 1 hour followed by incubation at 4°C overnight with primary antibodies (SYNGAP-1:2K, PA1-046, Thermofisher; b-Actin-1:5K, A2228, Sigma Aldrich). Membranes were washed thrice with TBST (0.1% Tween 20) followed by 1-hour incubation with HRP conjugated secondary antibodies (1:10K dilution) at RT. After washing the membranes three times with TBST, ECL (ECL-Prime western blotting system, GERPN2232, Sigma-Adrich, UK) was applied and digitally scanned using ImageQuant (ImageQuant LAS4000 scanner, GE healthcare and life Sciences). The density of individual bands was calculated using ImageJ (Version: 2.0.0). For SYNGAP levels, each value was normalized to β-actin and then to their control littermates. For pre/post-synaptic protein levels, each value was normalized to total protein and then to the average WT value.

### Open field

8 WT and 10 *Syngap^+/Δ-GAP^* male rats were acclimated to a holding room for at least 30 min before being individually placed in the corner of an empty grey painted wooden open arena (dimensions 100 x 100 x 50 cm, no bedding) evenly lit (avg. 40 lux on the floor). Spontaneous exploration was recorded for 20min/day on two consecutive days and activity measured using ANY-maze tracking and analysis software (Stoelting Co., IL, USA).

### Elevated plus maze

The elevated plus maze apparatus was raised 80 cm above the floor, made of dark plexiglass, and comprised of four arms (two open and two enclosed by 17 cm high walls). Arms were 70 cm long and 12 cm wide connected by a central square (dimensions 10 cm × 10 cm). 8 WT and 10 *Syngap^+/Δ-GAP^* male rats were acclimated to a holding room for at least 30 min before the start of the experiment. Rats were then placed individually at the central square of the apparatus facing an open arm and their spontaneous behaviour was recorded and tracked with ANY-maze tracking and analysis software (Stoelting Co., IL, USA) for 10 min.

### Rotarod

Rats were acclimated to the testing room for 60 min before being placed on individual lanes of a rotarod (Rotamex, Columbus Instruments, OH, USA) facing a white wall. Two trials of 90 sec each were performed at a constant speed of 4 rotations per minute (rpm) for baseline assessment of motor coordination. Rats were then left to rest for 30 min in their homecage. To assess motor learning, four trials of 90 sec each were performed, during which the rotarod speed started at 4 rpm and accelerated every 8 sec, until it reached 40 rpm. The above protocol was repeated for a total of 5 consecutive days. Latency to fall from the rotating drum was quantified through the Rotamex software and averaged across trials for analyses.

### Cued Fear Conditioning

12 WT and 11 *Syngap^+/Δ-GAP^* male rats were acclimated to a holding room and handled there for 5 min/day for two days before habituation to the testing context (a modified Coulbourne Instruments rat Habitest box dimensions 30 cm × 25 cm × 32 cm, containing a curved plastic black and white striped wall insert, smooth plastic grey floor, no electrified grid, scented with 70% ethanol by cleaning between trials) for two 5 min sessions on non-consecutive days (2 or 3 days apart). Conditioning followed on the day after the second habituation to the test context and was performed in a standard, unmodified Habitest rat box with aluminium wall inserts and electrified shock floor (Coulbourne H10-11R-TC-SF) cleaned with Distel™ disinfectant wipes between trials. Conditioning occurred over a 21 min period and consisted of a 3 min period to allow for exploration of the chamber followed by 6 pairings of a conditioned stimulus (CS) co-terminating with the unconditioned stimulus (US). The CS was a 10 sec blue flashing light (5 Hz 110 lux flashes, 50 / 50 duty cycle); the US was a 1 sec, 0.8 mA scrambled foot shock delivered through the bars of the floor; CS presentations started at 180, 360, 490, 770, 980, and 1280 sec into the training period. A separate control group (7 WT and 7 *Syngap^+/Δ-GAP^* males; CS-only) was exposed to 6 presentations of the CS alone in the same context. Before and after each session, rats rested in the holding room for at least 20 minutes. A video camera mounted above each context recorded the sessions. Percent time freezing was calculated for the first 9 sec of CS presentation during conditioning (when the footshock was absent). 24 hr after conditioning, retention of the conditioned response was tested. After rats were placed into the testing context, a 2 min period followed to allow for exploration, then twelve 30 sec long presentations of the CS, separated by 30 sec of no CS were given. An extinction index was calculated as the average freezing to CS 1-4 and CS 9-12/ total time freezing to CS 1-4 and 9-12; a modulation index was calculated as (time freezing_CS_ - time freezing_postCS_)/(time freezing_CS_ + time freezing_postCS_) for average freezing to CS 1-3 (early) and CS 9-11 (late).

### Spatial reference memory water maze

8 WT and 9 *Syngap^+/Δ-GAP^* male rats were trained in three stages in a 2 m diameter water maze containing a 10 cm escape platform. Water was made opaque using liquid latex (Palace chemicals, Liverpool, UK) and kept at a temperature of at 18-20 *C. First, rats were trained for 2 days on the visible platform version of the water maze (4 trials / day, 15 min ITI, extra-maze cues obscured by a white curtain). In the second stage, the curtain was removed and wallmounted extra-maze cues approx 1 m from the edge of the pool were visible. Rats received one daily hidden-platform training session (4 trials / day, 15 min ITI) for 6 consecutive days. Reinforced probe trials were given on the first trial on the 3^rd^ and 6^th^ day of training, followed by three standard training trials separated by 15 min ITI. During reinforced probe trials, an ‘Atlantis’ platform (Spooner *et al*., 1994) was used, which is submerged to a depth the animals cannot reach during the 1 min of the probe trial, but then automatically raises to the same depth the platform has during the training trials, i.e., 4 cm below the water surface. Each trial lasted a maximum of 2 min; rats failing to escape were guided to the platform. All rats remained on the platform for 15 sec before removal from the pool. The final (reversal) stage of the protocol started the following day and was identical to the second, with the exception that the platform was relocated to the opposite side of the pool. Platform locations were counterbalanced across genotypes. Release location was pseudorandomised for each trial and counterbalanced for genotypes across all days. During the ITI rats were dried with a towel and were returned to a holding cage (identical to their homecage), which was placed on a heating pad with monitored temperature. A video camera mounted above the pool recorded the sessions through WaterMaze software to obtain swim paths, path lengths, and swim speed. Data was averaged across trials for analyses, with the exception of data recorded during probe trials.

### Spontaneous object exploration tasks

5 WT and 8 *Syngap^+/Δ-GAP^* male rats underwent object recognition (OR), object-place recognition (OPR), object-context recognition (OCR) and object-place-context recognition (OPCR) testing as previously described (Langston & Wood, 2010; Till *et al*., 2015). Briefly, rats were tested in a rectangular testing box (dimensions 60 cm x 40 cm x 50 cm) with removable walls and floor inserts that could change into two context configurations. In context 1, white textured wallpaper and laminate floor were used. Blue wood laminate walls and a black rubber floor were used for context 2. After 5 consecutive days of habituation to the boxes (5 min / day), rats received 2 trials (one/day) on each of the four tasks, consisting of a 3 min sampling phase(s), a 2 min ITI, and a 3 min test phase. A video camera above the box recorded the sessions for subsequent scoring of time exploring, by quantifying time rats spent sniffing the objects. If rats did not reach a 5 sec minimum of exploration for both objects or a 15 sec minimum of total object exploration during the sample phase, or did not reach a minimum of 15 sec of total object exploration in the test phase, their measures on that task were excluded from analysis as it cannot be confirmed they spent enough time exploring to learn/discriminate. For each test phase, a discrimination index *d* was calculated as follows: [(time exploring novelty—time exploring familiarity)/(sum time exploring)]. To determine whether animals prefer the novelty, observed index *d* was compared against chance performance (score of *d* = 0.0) using a two-tailed one-sample *t-test*. Values significantly above *d* = 0.0 indicate preference for novelty. During ITIs, rats were placed in a covered plastic holding bucket containing sawdust. All objects, locations, and/or contexts were counterbalanced for trial and genotypes.

### Three chamber task

Rats were habituated for 3 consecutive days to the testing apparatus: a plexiglass rectangular box (dimensions 150 cm x 50 cm x 30 cm), divided into three chambers; left and right chambers (60 cm x 50 cm each) communicated to the centre chamber (30 cm x 50 cm) via removable doors. After 2 min exploration of the central chamber, doors were opened to enable the test rat to explore the entire arena for 10 minutes. For the first habituation session (day1, H_1_), all three chambers were empty, whereas for consecutive habituations days 2-4 (H_2_-H_4_) each outer chamber contained one wire cage. After the last habituation session, the test rat was removed from the apparatus and placed in a covered plastic holding bucket containing sawdust for 5 min before phase 1 began. Rats were either tested in a social interaction or a social preference task. 12 WT (8 male, 4 female) and 14 *Syngap^+/Δ-GAP^* (7 male, 7 female) rats were used to assess social interaction. After exploring the central chamber for 2 min, the doors were raised, and the test rat was free to explore the entire arena for 10 min; one wire cage was left empty and the other contained a non-familiar wild-type Long-Evans rat of the same-sex and similar age. Data from males and females were pooled for this analysis as both sexes showed the same level of preference for the rat over the empty (males: effect of stimulus F_(1,13)_=36.39, *p*<0.0001; females: effect of stimulus F_(1,9)_=51.55, *p*<0.0001). A separate cohort of rats (10 WT and 10 *Syngap^+/Δ-GAP^* males) were used to assess social preference. In this task configuration, one wire cage contained a non-familiar rat while the other cage contained a novel object. Rats used as social stimuli were habituated to being restrained in the wire cages for at least 3 days prior to the start of the experiment, by simulating the entire procedure with WT Long-Evans rats (not used as testing animals). A video camera above the apparatus recorded the sessions for subsequent scoring of time in close interaction, by quantifying time rats spent actively sniffing, and time in chamber through ANY-maze tracking and analysis software (Stoelting Co., IL, USA). Sociability index was calculated as follows [(time exploring rat—time exploring empty)/(sum time exploring)], social preference index was calculated as follows [(time exploring rat—time exploring object)/(sum time exploring)]. To determine whether animals prefer the social stimulus, the observed index was compared against chance performance (theoretical *u* = 0.0) using a two-tailed one-sample t-test. Values significantly above 0.0 indicate preference for social over non-social. The apparatus was thoroughly cleaned with baby wipes and 70% ethanol between trials. During ITIs, rats were placed in a covered plastic holding bucket containing sawdust. All experiments were counterbalanced across conditions: location, ID of stimulus rat, genotype and sex.

### EEG with 32-channel skull-surface grid probe

12 WT and 12 *Syngap^+/Δ-GAP^* male rats were anaesthetised and prepared for stereotaxic surgery. Two craniotomies were drilled for bilateral anchor screw placement (+4.0 mm AP, ± 0.5 mm ML) and one for ground screw implantation (−11.5 mm AP, 0.5 mm ML), according to the frontal and caudal edges of the EEG array probe (H32-EEG – NeuroNexus, MI, USA). The EEG probe was placed on the skull with its cross symbol aligned over bregma. The ground electrode and screw were connected, the implant was covered with dental cement, and animals were allowed to recover for a minimum of 1 week post-surgery. Prior to recording, rats were habituated to the room. On recording days, up to 4 rats, in their individual home cages, were placed concurrently inside a 1 x 1 m faraday cage. 6 hour EEG recordings were acquired with an Open Ephys (Siegle et al., 2017) acquisition system (OEPS, Portugal), through a 32-channel recording headstage amplifier linked to an accelerometer (RHD2132 Intantech, USA), at a sampling rate of 1 KHz.

### Manual detection of SWD

For off-line visual seizure scoring, 6 EEG channels were selected from each recording and analysed on a custom-designed interphase using Igor Pro V6.3 (Wavemetrics, OR, USA). After identifying the presence of SWD events, as well as wake and sleep epochs, the data was visually scored in successive 0.2 sec epochs by an observer blinded to genotypes SWDs with an inter-SWD interval shorter than 1 sec were considered as one, while individual SWDs shorter than 0.8 sec were discarded.

### Automatic detection of SWD

Spectral analysis revealed that visually scored SWDs behave as a high energy echo of a fundamental frequency (f0) located on the 5-10 Hz theta band (7.7 ± 0.1 on both genotypes), that resonates in several periodic harmonics across the frequency spectrum. (**Supplementary Figure 5A**). This oscillating spectral structure of SWDs resembles a periodic waveform, allowing their automatic identification through a cepstral analysis approach (Childers et al., 1977) by searching for a high amplitude peak located on a frequency band of interest. We applied an automated SWD seizure detection algorithm to voltage traces from the EEG grid electrode lead overlaid approximately on S1 (right hemisphere, AP −3.0 mm and ML 2.8 mm from bregma), as, by visual assessment, was the channel most frequently associated with high amplitude SWDs across animals. After deconvolving the raw signal using a Fast Fourier Transform (number of tapers =5), a logarithm was applied to obtain the magnitude. The signal could then be treated as semi-periodic so that the inverse Fast Fourier Transform could be applied to obtain the cepstrum and reveal the period of the fundamental frequency (f0) as a spike in a pseudo-time domain frequency (**Supplementary Figure 5B**). After obtaining the cepstrum for the entire EEG recording (in sliding windows of 0.2 sec), peak power cepstrum values within the relevant frequency range (5-10 Hz) were identified and normalized by their absolute maximum. The resulting vector was transformed into z-scores to homogenise possible power differences between recordings that could distort seizure threshold identification. A threshold of ≥ 2.2 x 10^-5^ standard deviations was set by comparing the values of visually scored seizures against other high magnitude noise that resulted in false positives (**Supplementary Figure 5C**). 0.2 sec time windows were time-stamped as seizures when z-scored peak cepstral power in the theta band was greater than or equal to the established standard deviation threshold (**Supplementary Figure 5D**). As in visual analysis, time-stamped SWDs with an inter-SWD interval shorter than 1 sec were considered as one, while individual SWDs shorter than 0.8 sec were discarded. For validation, the results from the automated method were compared against the visual analysis and show that over a recording period of 6 hrs 100% of the visually counted SWDs were accurately detected, as confirmed by a non-significant difference between the two methods in the number of SWDs detected (paired *t-test*; t_(10)_= 1.624, *p*=0.135; **Supplementary Figure 5E**). Automatically detected SWDs were also compared between genotypes, obtaining a significant statistical difference equivalent to that found by visual scoring (Mann-Whitney U test, visual detection: U=23, *p*=0.002; automatic detection: U=25, *p*=0.003, **Supplementary Figure 5F and 5G** respectively). The code used for analysis of this section in the study is freely available via GitHub repository (https://github.com/Gonzalez-Sulser-Team/SWD-Automatic-Identification).

### Pharmacological suppression of SWD

2-hour EEG recordings were performed daily over 5 consecutive days (see Figure **7A** for treatment timeline). Briefly, animals received no treatment on days 1, 3 or 5. On days 2 and 4, recordings were made starting 1 hour after animals received a single treatment of either ETX (100 mg/mL, Sigma-Aldrich) or 0.9% saline (vehicle) with a volume dose of 1mL/kg delivered by intraperitoneal injection. Animals were counterbalanced for whether drug treatment was received before saline or *vice versa*. SWDs were quantified using the automatic seizure detection method described above by a researcher blinded to genotype and drug treatment used on each day. A seizure suppression index was calculated over the two hours of recording as follows: ((SWD_treated_ - SWD_pre-treated_)/ (SWD_treated_ + SWD_pre-treated_)). A negative value indicates fewer seizures than pre-treatment, whereas a positive value indicates more seizures than pre-treatment.

### Statistical analysis

Unless otherwise stated, error bars in all graphs indicate standard error of the mean (sem) and all statistical tests were two-tailed. Unless otherwise stated, mean, median, standard error and statistics were calculated across animals. Where 3-way ANOVAs were performed (i.e figure 2), we used a mixed-effects restricted maximum likelihood (REML) model with genotype as a matching factor. This was because group sizes were different for *CSonly* and *CS-US* paired. Exact *p* values are reported within the text. All the statistical tests performed can be found in **Supplementary Table 1**. In the figures, asterisks denote significant results for alpha set at 0.05. \**p*<0.05, \*\**p*<0.01, \*\*\**p*<0.001, \*\*\*\**p*<0.0001. Diamonds illustrate above chance performance with *p*<0.05.

## Acknowledgements

The authors would like to thank all members of the Kind laboratory for their helpful discussions. We thank our funders: the Simons Initiative for the Developing Brain, Patrick Wild Centre, Medical Research Council UK (MR/P006213/1), FRAXA Foundation, Department of Biotechnology India, Wadhwani Foundation, the Shirley Foundation, The Wellcome Trust (204954) and RS Macdonald Trust.

## Author contributions

D.K., S.M.T., M.J., M.A.C., S.C., A.G.S., E.R.W., O.H., P.C.K. designed the experiments. D.K., S.M.T., I.B.P., T.C.W., M.N., D.A., S.T., V.K., J.S., N.A., L.M. performed the experiments. D.K., S.M.T., I.B.P., T.C.W., S.N., S.T., V.K., N.P. analysed the data. D.K., S.M.T., and P.C.K. wrote the manuscript with input from all authors.

## Supplementary Methods and Figures

### EEG with implanted screw electrodes

Rats were anaesthetised and prepared for stereotaxic surgery. Craniotomies were drilled and a single recording screw positioned at each of the following coordinates relative to bregma: +7.56 mm AP, 1 mm ML (olfactory bulb), +2.16 mm AP, 3 mm ML (motor cortex), −3.24 mm AP, 2.5 mm ML (parietal association cortex). A screw positioned over the cerebellum served as reference/ground (−12 mm AP, 0 mm ML). Recording screws were implanted unilaterally and then connected to an electronic interface board (EIB 16, Neuralynx). The incision was then closed using surgical sutures (Ethicon, Henry Schein, UK) and rats were left to recover for a minimum of 1 week post-surgery. Recordings were made via a 16 channel digitising headstage (C3334, Intan Technologies, USA) in the same system as described in main Methods. LFP signals were bandpass-filtered from 0.1 - 600 Hz and sampled at 2 kHz in OpenEphys software. Video recordings were made using Freeze Frame software (15 frame per sec; Actimetrics) synchronised with electrophysiological signals via TTL pulses.

### Visual stimulation during EEG recordings

EEG recordings were made from implanted 6 WT and 6 *Syngap^+/Δ-GAP^* male rats within a 35 x 20 x 40 cm plastic cage positioned within a sound attenuating chamber. Rats were given 2 min to explore the context prior to presentation of a 10 s visual stimulus (5 Hz 110 lux flashes, 50 / 50 duty cycle). This was followed by a post-stimulus time of >1 min with recordings maintained throughout. SWD events were manually identified. Their total number was quantified and their timing relative to the onset of the visual stimulation was calculated in 10 sec bins. Neuroexplorer software (Nex Technologies, CO, USA) was used to generate spectrograms (0.4 sec shifting window, 50% shift overlap, time-bandwidth product = 3, number of tapers = 5).

### Olfactory habituation-dishabituation task

Rats are transferred to an empty cage, similar to their home cage, and a series of odour-infused cotton swabs are presented for 2 min each, with a 1 min ITI (adapted from (Yang & Crawley, 2009)). Rats are acclimatised to the testing environment a day prior to the experiment, by being placed in an empty cage with a cotton swab infused with ddH_2_O. The odour order is as follows:

> *ddH_2_O, ddH_2_O, non-social odour 1.1, non-social odour 1.2, non-social odour 1.3, nonsocial odour 2.1, non-social odour 2.2, non-social odour 2.3, social odour 1.1, social odour 1.2, social odour 1.3, social odour 2.1, social odour 2.2, social odour 2.3, ddH_2_O*

Banana extract (1:1000 diluted in ddH_2_O; Foodie Flavors™) and almond extract (1:1000; Foodie Flavors™) were used as non-social odours. Swabs of the bedding surface of home cages of 4 group-housed adult rats (sex-matched) were used as social odours. Odours were counterbalanced for order of exposure.

**Supplementary Table 1.**
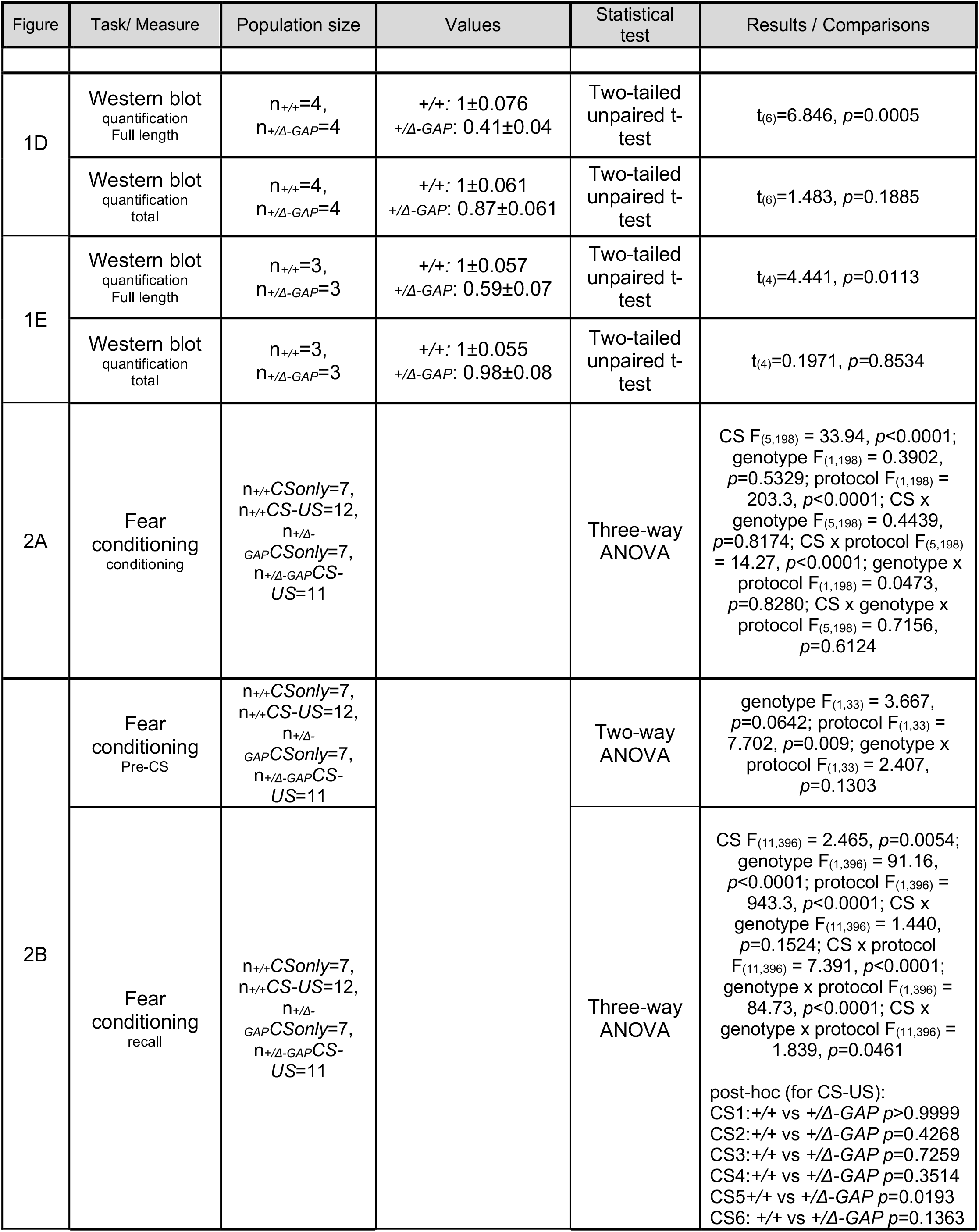
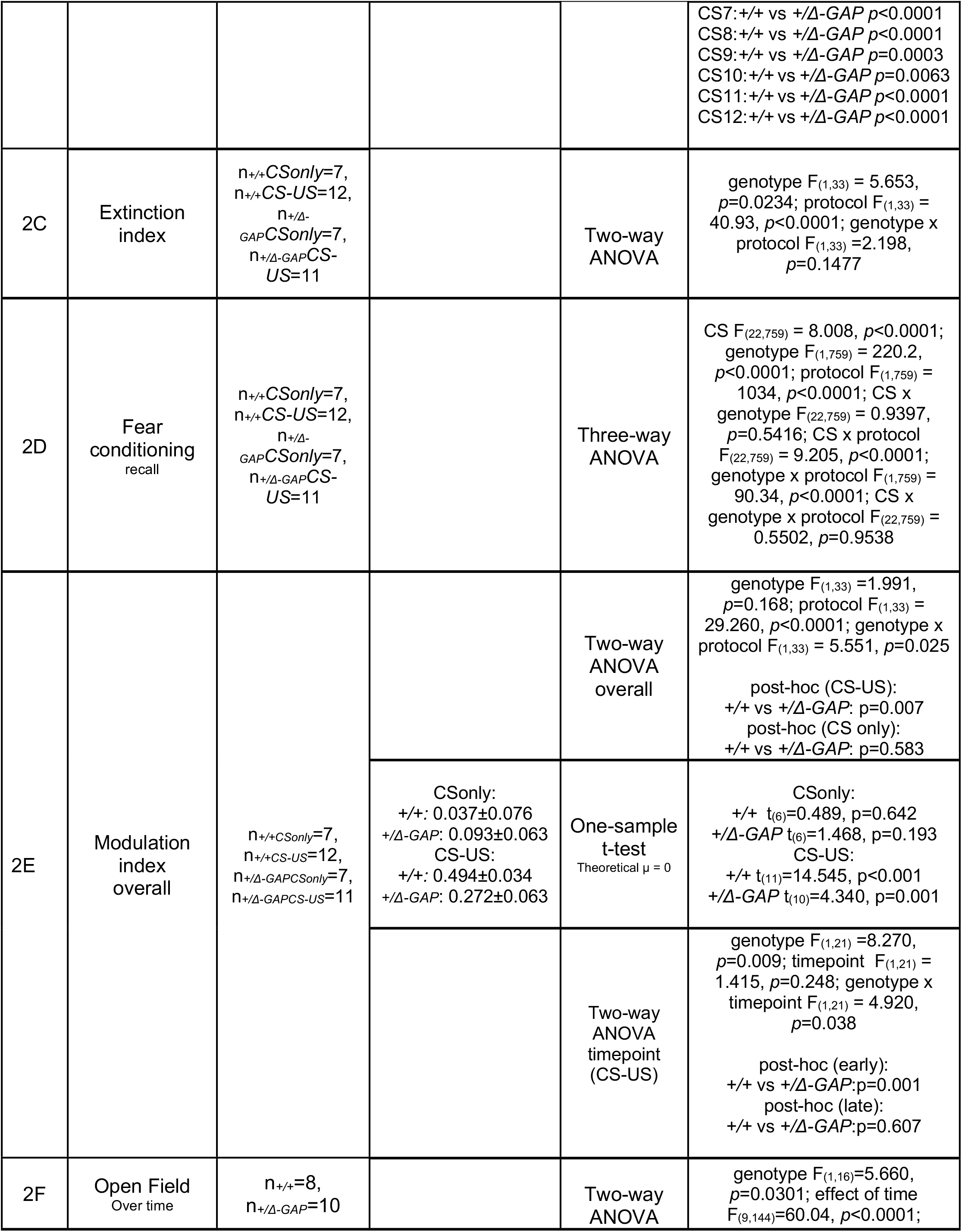
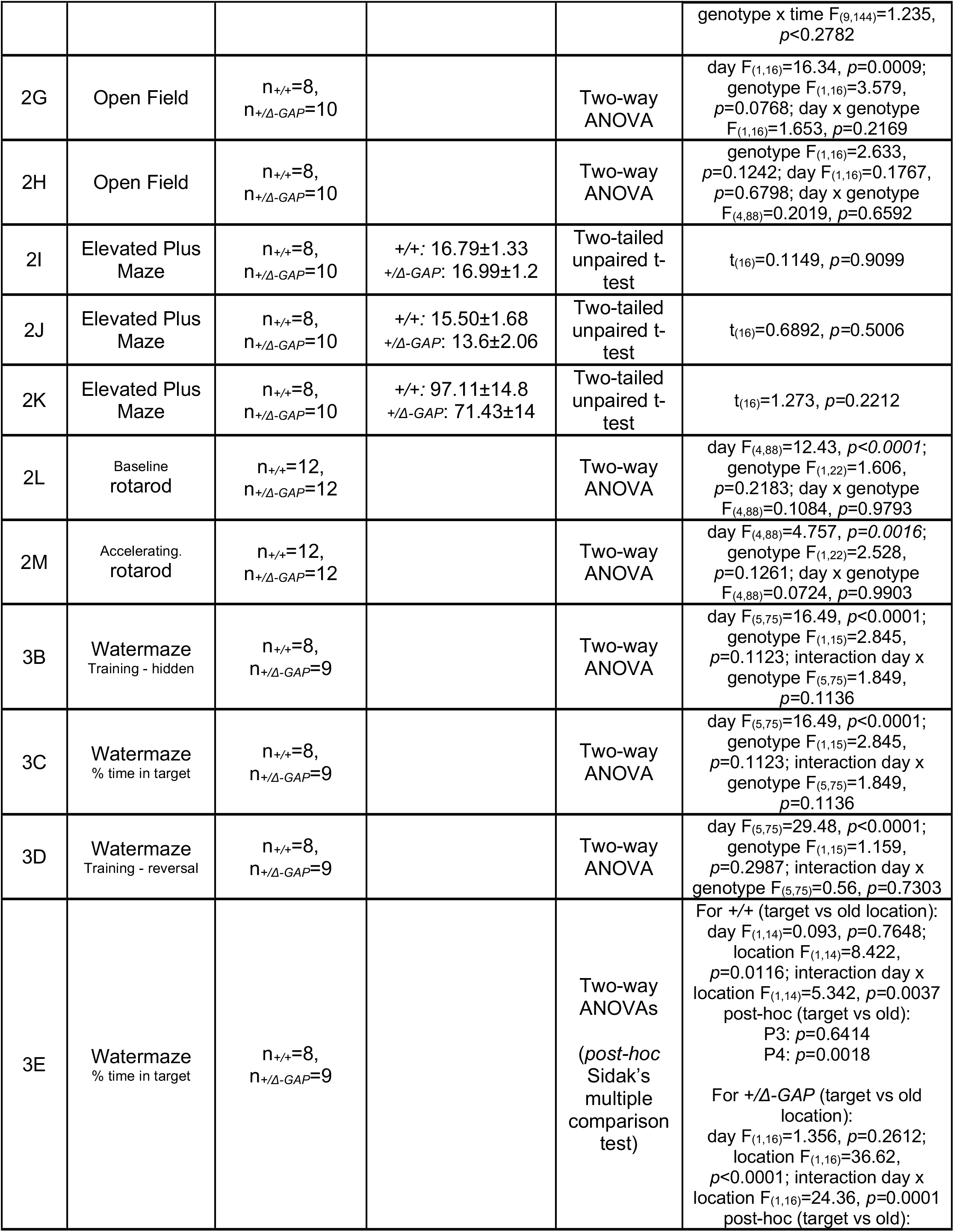
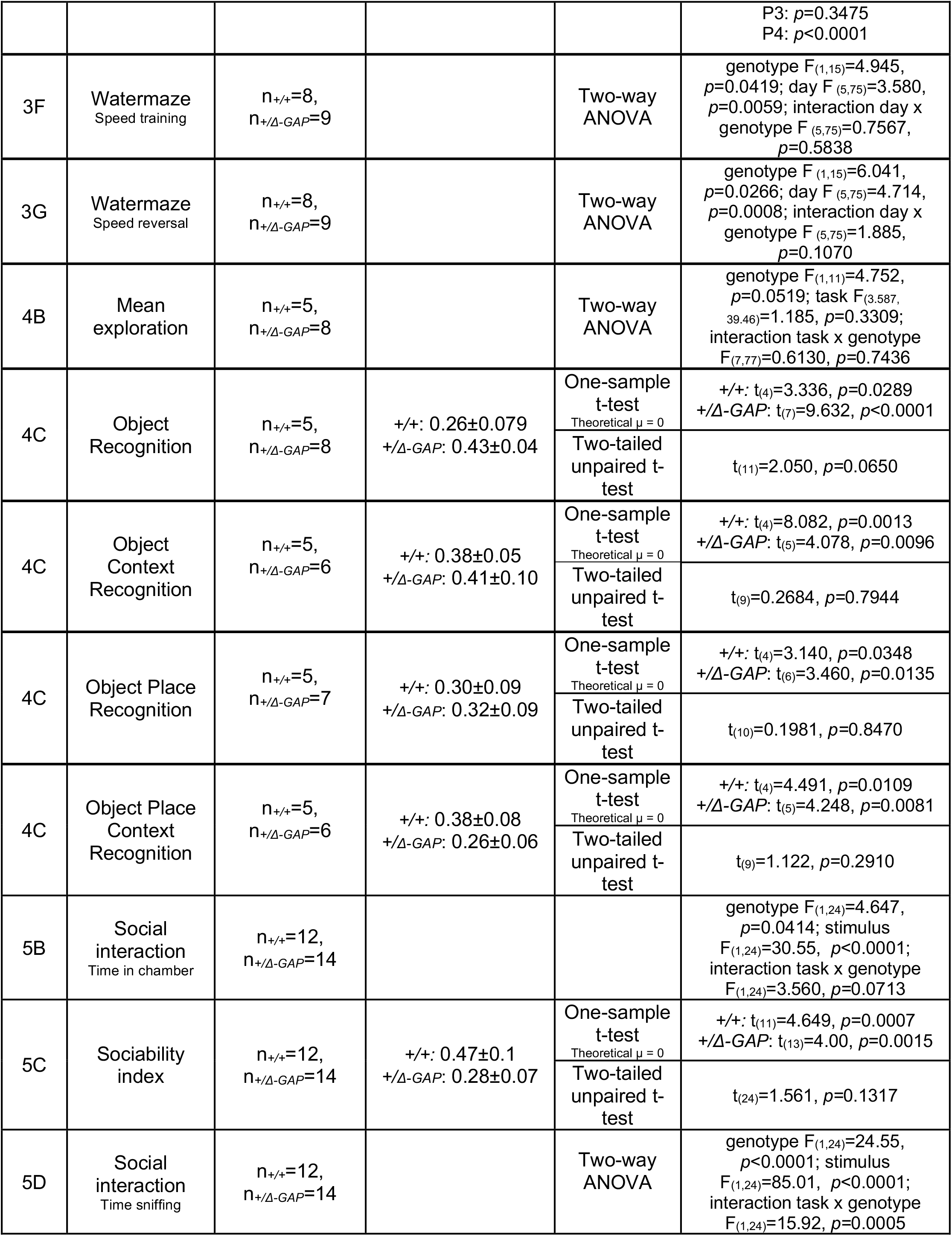
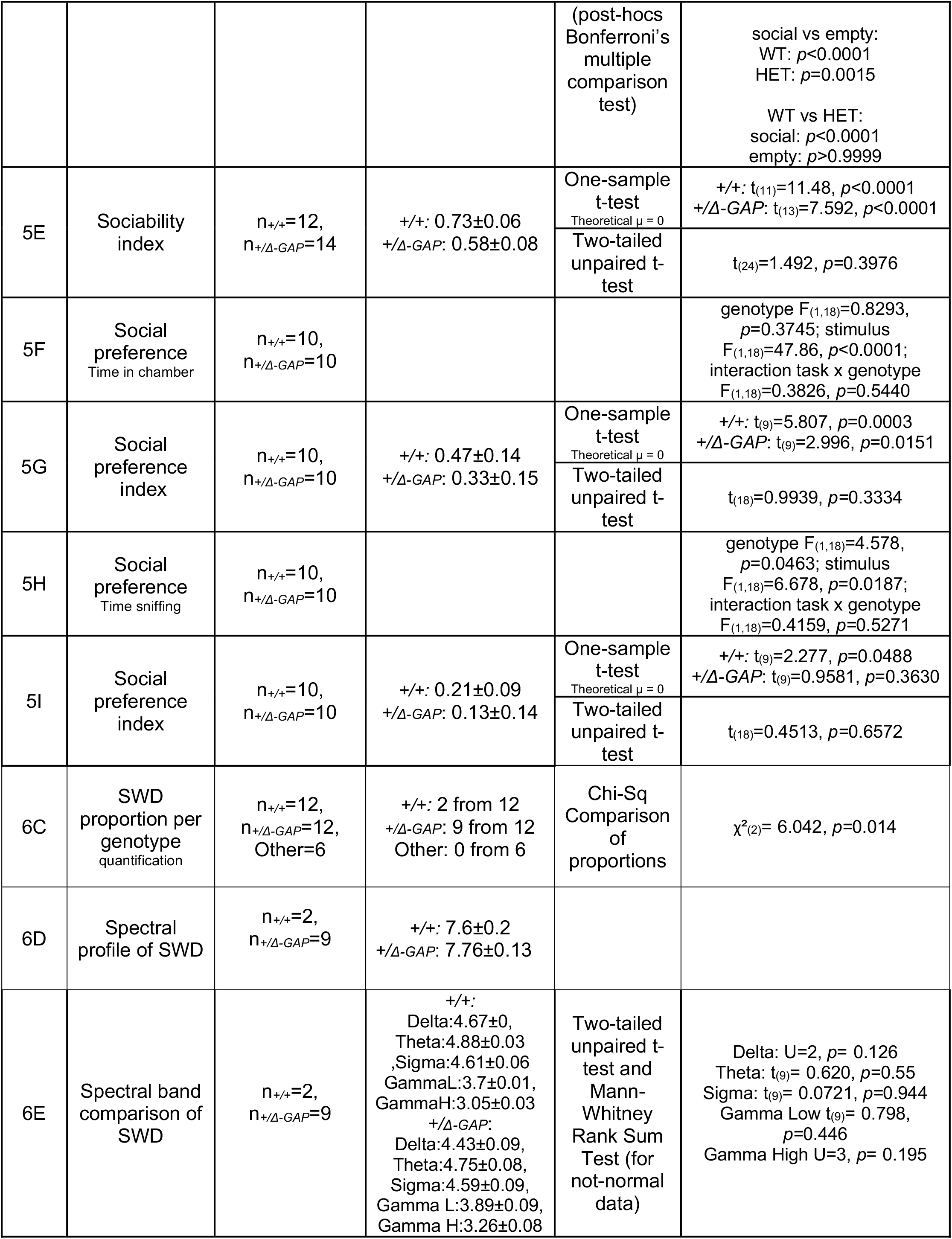
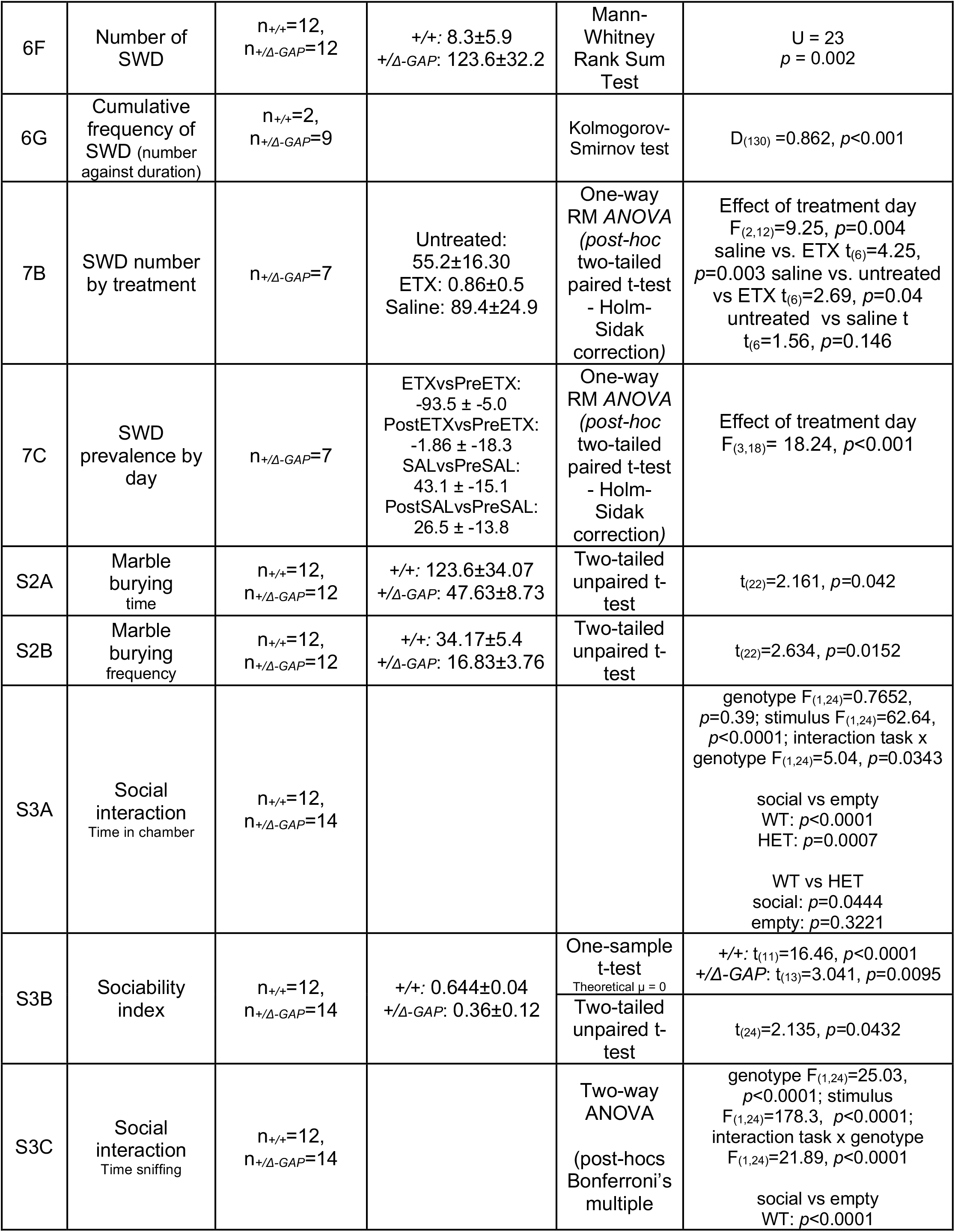
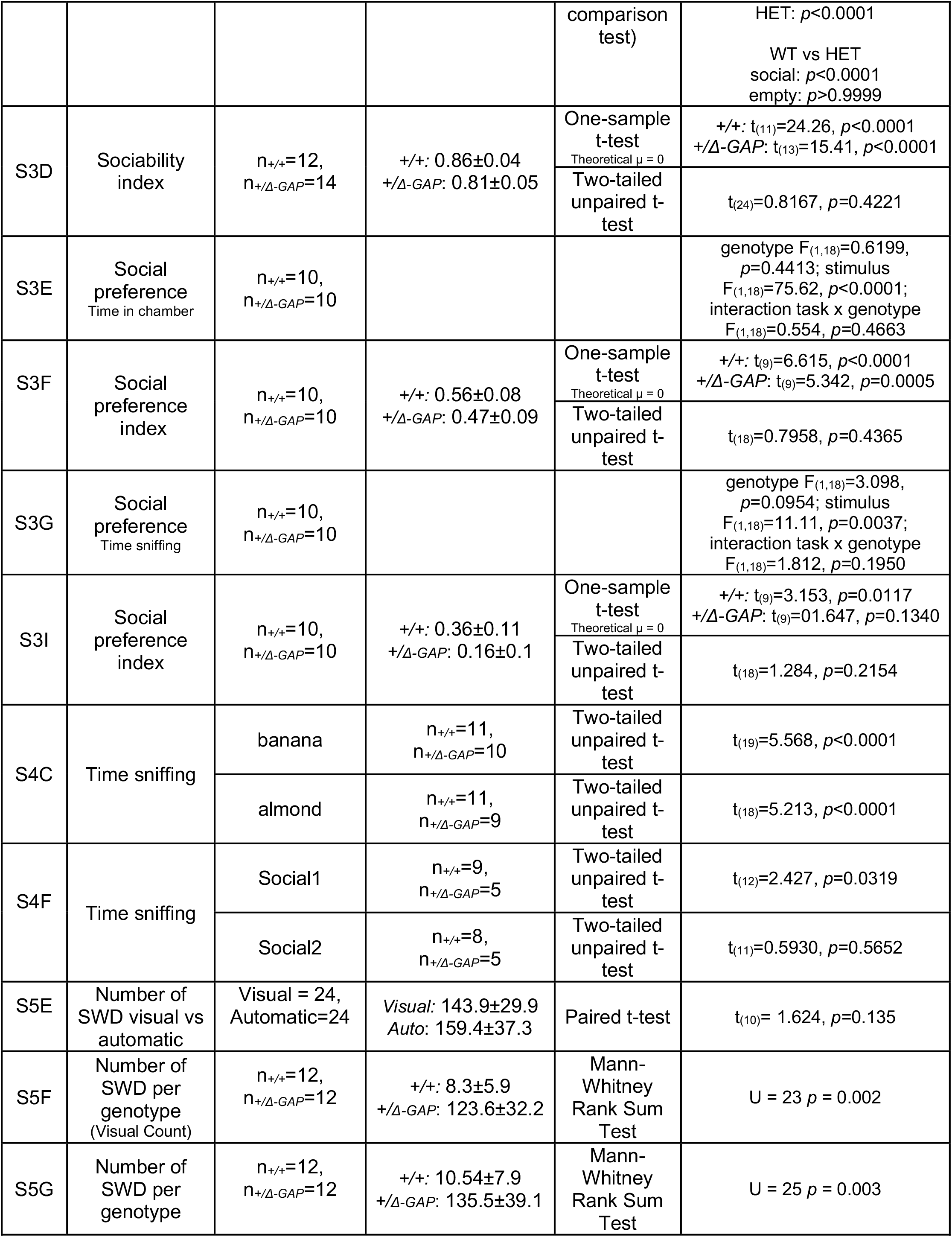
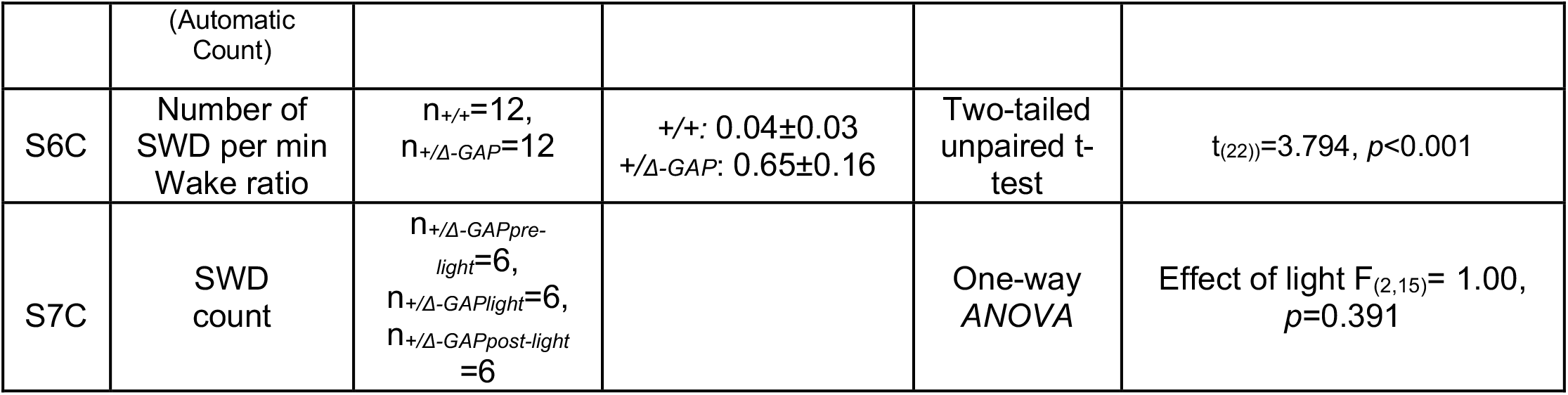
Summary of data and exact p-values related to Figures. Applicable figure panels are listed followed by the relevant measure (task) for which data is reported. Data values are given in mean ± SE. Statistical tests (t-tests, ANOVAs) are then followed by descriptive statistics and exact *p*-values for results and comparisons made for each relevant measure. All one-sample t-tests are compared against chance level (theoretical mean of 0.0). *+/+* for *Syngap^+/+^*, *+/Δ-GAP* for *Syngap^+/Δ-GAP^*. See Methods.

**Supplementary Figure 1.**
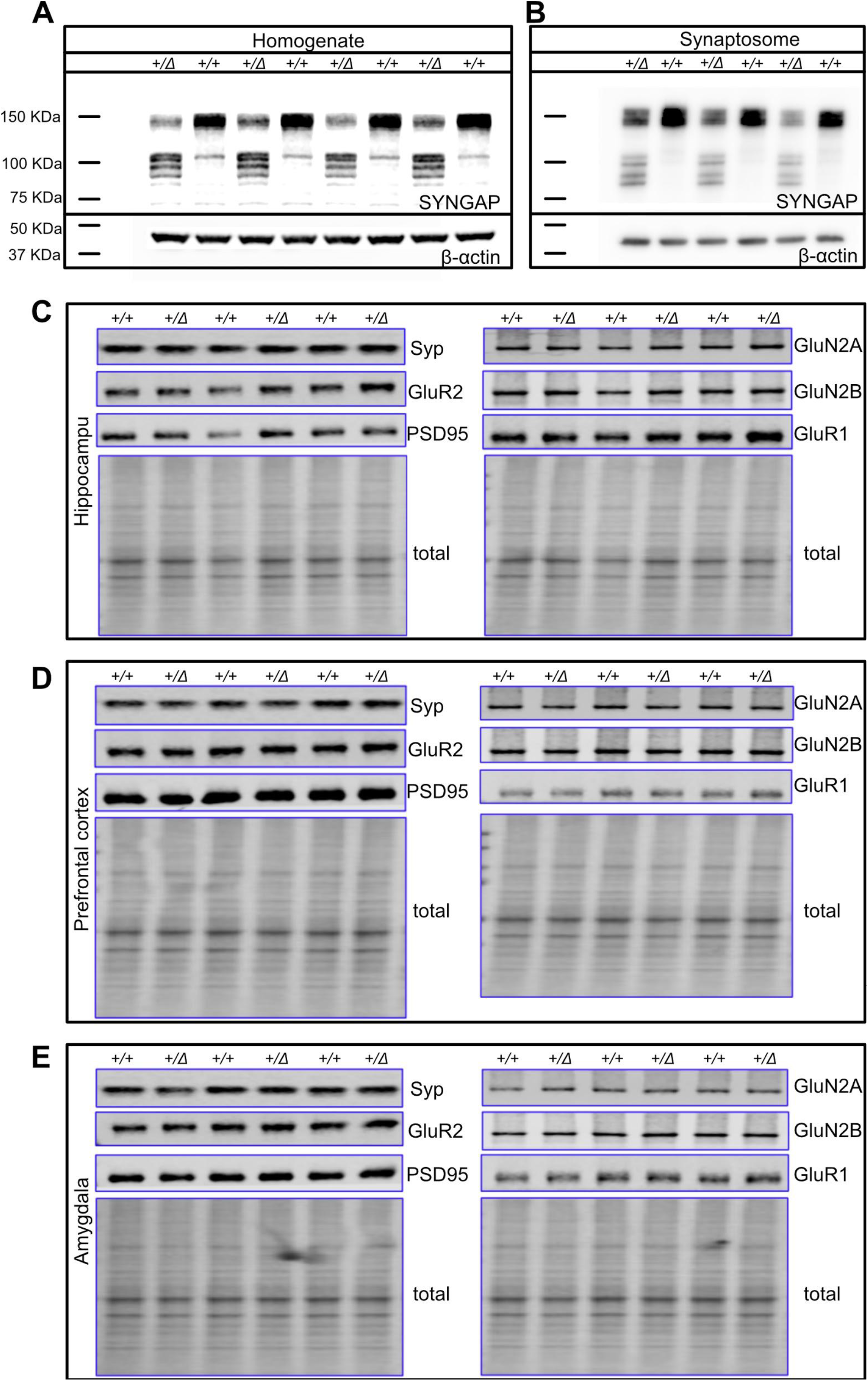
Full-lengthSYNGAP levels are reduced in *Syngap^+/Δ-GAP^* rats. Western blots of (A) homogenates (n_+/+_ = 4, *n_+/Δ-GAP_* = 4) and (B) synaptosomes (n_+/+_ = 3, *n_+/Δ-GAP_* = 3) from adult rat hippocampus confirm that full-length endogenous SYNGAP protein (~150kDa) is located in synapses; additional bands in the molecular weight range predicted for mutant SYNGAP isoforms are detected in both homogenates and synaptosomes from *Syngap^+/Δ-GAP^* rats. Western blots of purified synaptosomes (n_+/+_ = 3, *n_+/Δ-GAP_* = 3) from adult rat hippocampus (C), prefrontal cortex (D), and amygdala (E) to quantify the level of several proteins associated with pre- and post-synaptic function. Syp; synaptophysin, GluR1; Glutamate receptor 1, GluR2; Glutamate receptor 2, PSD95; Postsynaptic density protein 95, GluN2A; NMDA receptor subtype 2A, GluN2B; NMDA receptor subtype 2B, +/Δ; +/Δ-GAP. Tissue from 3 animals was pooled for each sample.

**Supplementary Figure 2.**
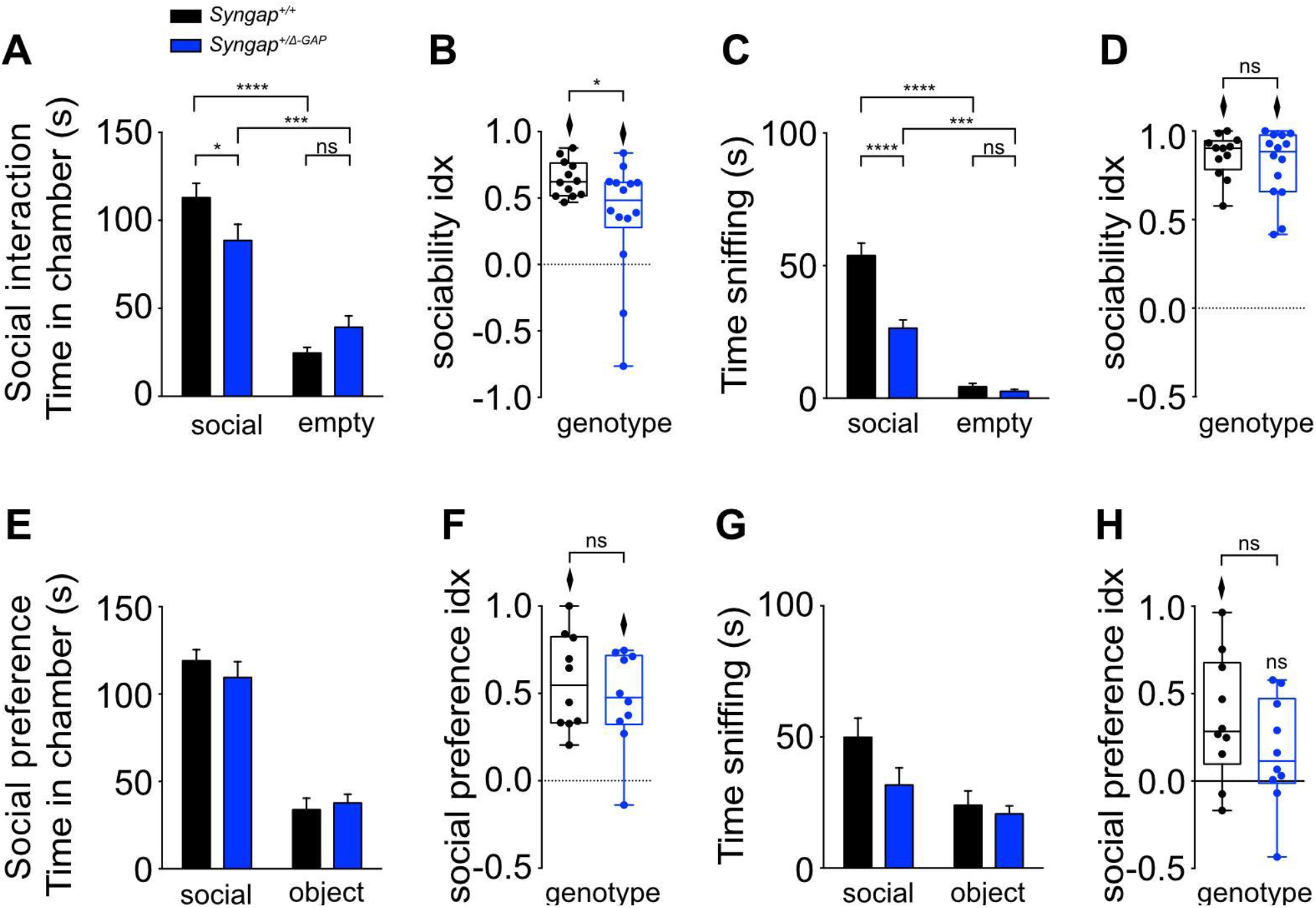
Social behaviour data for 0-180 seconds. In the social interaction task, time in chamber (A) and sociability index (B) indicate WT and *Syngap^+/Δ-GAP^* rats show a preference for spending time in the chamber containing a caged social stimulus compared to the chamber containing an empty wire cage. WT rats were more reliably spending time in the social chamber than *Syngap^+/Δ-GAP^* rats. (C) Time actively exploring (sniffing) and (D) sociability index for active exploration suggest that both genotypes explore the social stimulus more (n_+/+_ = 12, *n_+/Δ-GAP_* = 14). In the social preference task, time in chamber (E) and social preference index (F) indicate WT and *Syngap^+/Δ-GAP^* rats spend significantly more time in the chamber containing a caged social stimulus compared to the chamber containing a novel object. (G, H) *Syngap^+/Δ-GAP^* rats do not show preference for actively exploring the social stimulus over the object (n_+/+_ = 10, *n_+/Δ-GAP_* = 10). Diamonds illustrate above chance performance (*p*<0.05).

**Supplementary Figure 3.**
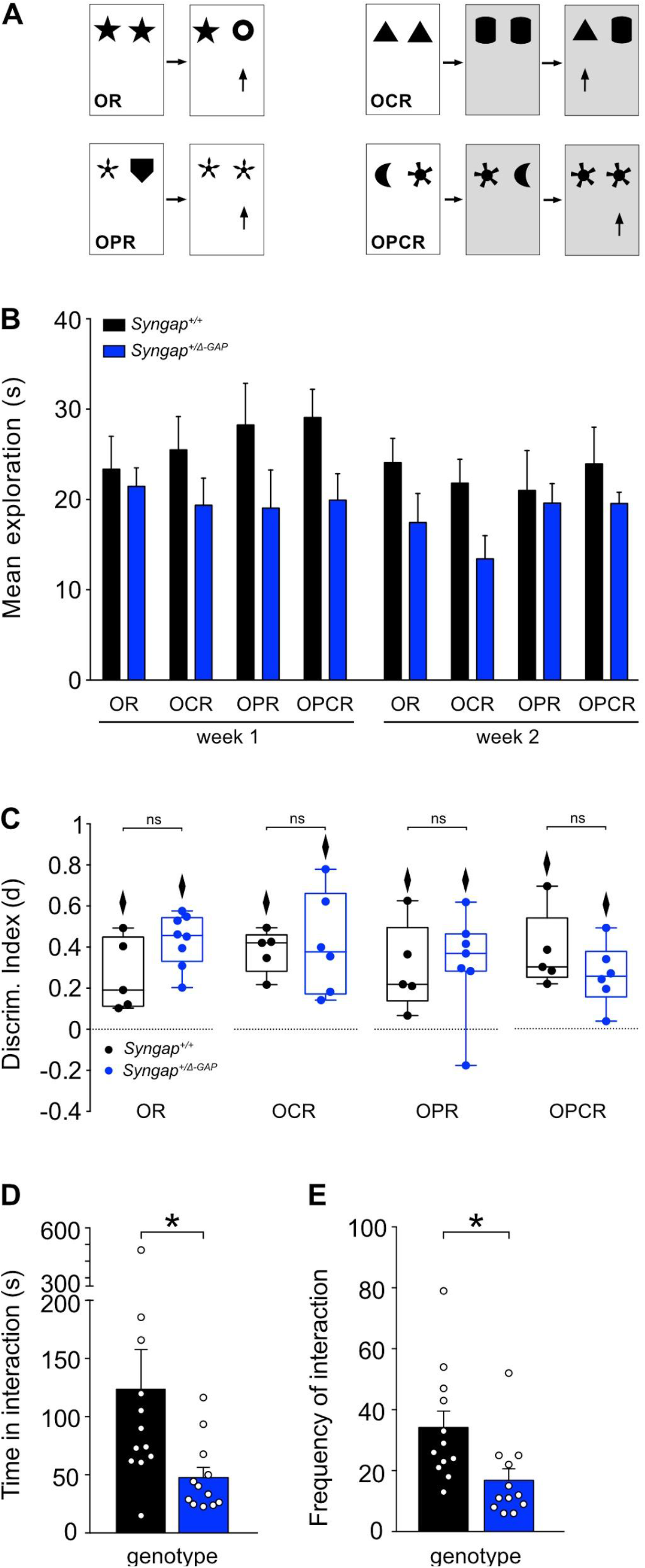
Associative recognition memory remains unaffected after heterozygous deletion of the C2/GAP domain in SYNGAP. (A) Schematic illustration of spontaneous exploration tasks for object (OR), object-context (OCR), object-place (OPR) and object-place-context (OPCR) novelty recognition. (B) *Syngap^+/Δ-GAP^* rats spend less time exploring during the first exposure of the objects in different configurations. (C) Both WT and *Syngap^+/Δ-GAP^* rats that reach exploration criterion (see methods) exhibit short term memory for all four tasks, as measured by above chance performance (illustrated by diamonds for *p*<0.05). In the marble burying task, *Syngap^+/Δ-GAP^* rats (D) spent significantly less time interacting with marbles and (E) their interactions were less frequent compared to WT littermates. *mean* ± SE is noted.

**Supplementary Figure 4.**
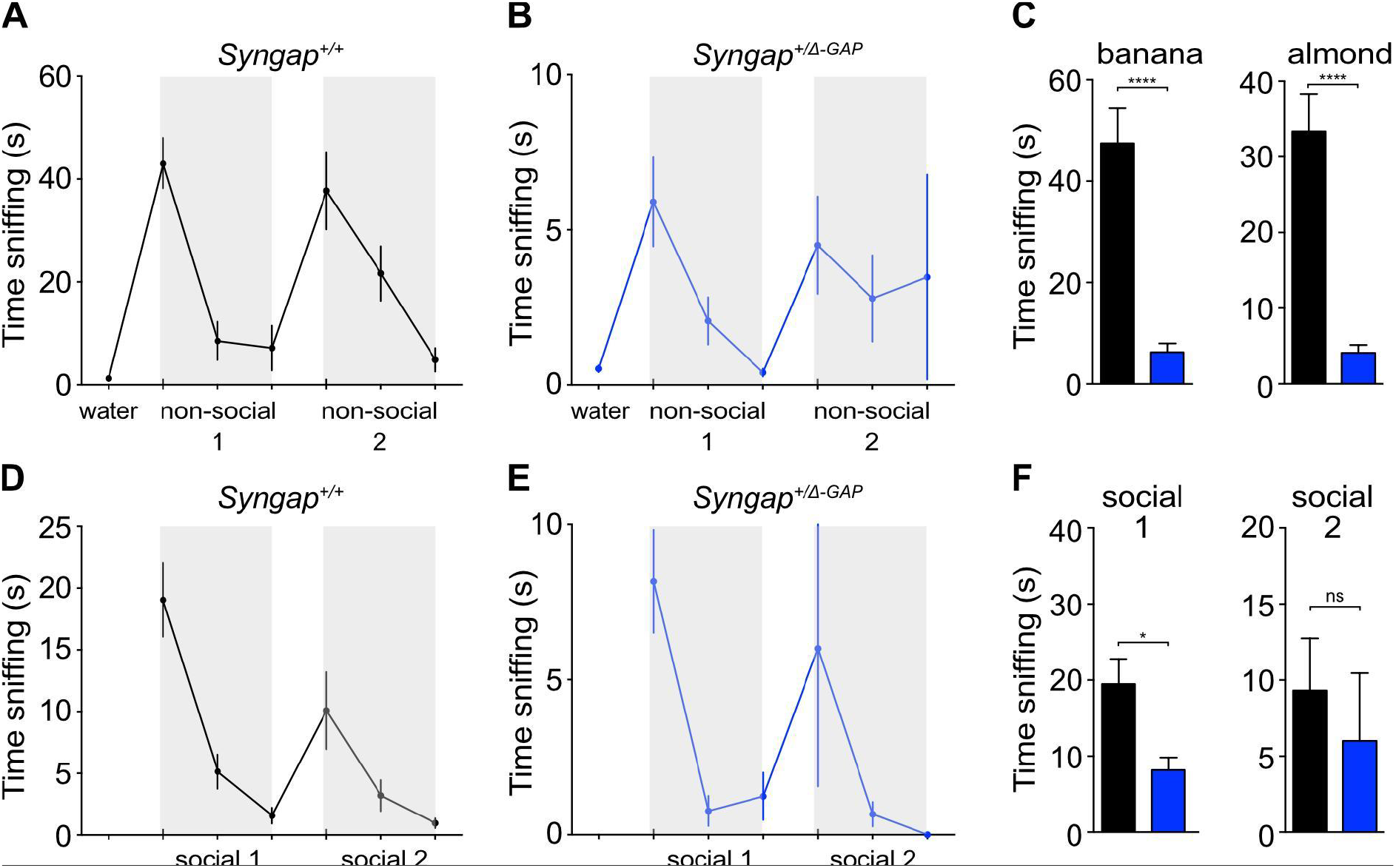
*Syngap^+/Δ-GAP^* rats can detect and discriminate odours. The average time WT (A) and *Syngap^+/Δ-GAP^* (B) rats spent investigating non-social odours habituates over three consectutive presentations of an individual non-social odour; a relative increase in time spent investigating a new non-social odour indicates olfactory discrimination. (C) *Syngap^+/Δ-GAP^* rats spent less time overall investigating each non-social odour than WT rats. WT (D) and *Syngap^+/Δ-GAP^* (E) rats reduce the amount of time spent investigating social odours over three consectutive presentations; a relative increase in time spent investigating a new social odour indicates olfactory discrimination. (F) *Syngap^+/Δ-GAP^* rats spent less time overall investigating each social odour than WT rats. Colour change indicates odour change. *mean* ± SE is noted.

**Supplementary Figure 5.**
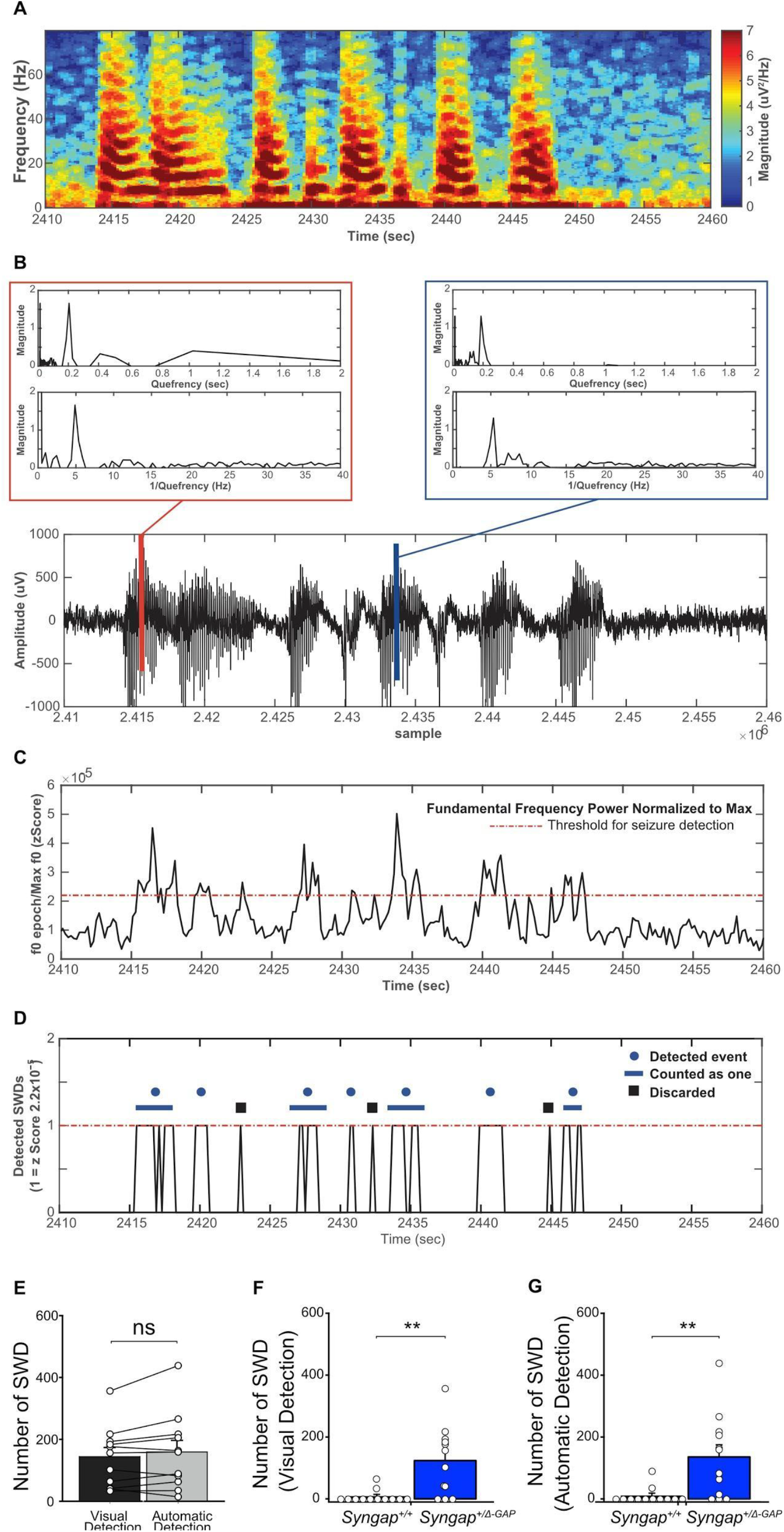
Automatic Detection of SWD. (A) Example spectrogram of SWD. (B) Raw EEG trace corresponding to time interval of the spectrogram in A (bottom), with two selected SWD epochs (0.2 sec each) marked with blue and red vertical lines. Inserts (top) show respective cepstral power analysis showing the fundamental frequency (f0) peak on the pseudo-time domain (top inset) and pseudo-frequency domain (bottom inset). (C) Peak cepstral power on theta band range (5-10 Hz) calculated in 0.2 sec epochs, normalized to its absolute maximum value, and transformed into z-scores. Threshold for detecting SWD events marked with dashed red line. (D) Detected SWDs transformed into zeros (below threshold) or ones (threshold or over) on the time interval shown in A and B. Events counted as one (< 1 s between events) are marked by blue lines, whereas black squares designate discarded events (length < 0.8 s). (E) Total number of SWD detected by visual (black bar) and automatic counting (gray bar) is comparable. (F, G) Genotype comparison of visually detected and automatically detected SWDs shows a significantly increased number of SWD events in *Syngap^+/Δ-GAP^* rats with both methods. (n_+/+_ = 12, *n_+/Δ-GAP_* = 12). *mean* ± SE is noted.

**Supplementary Figure 6.**
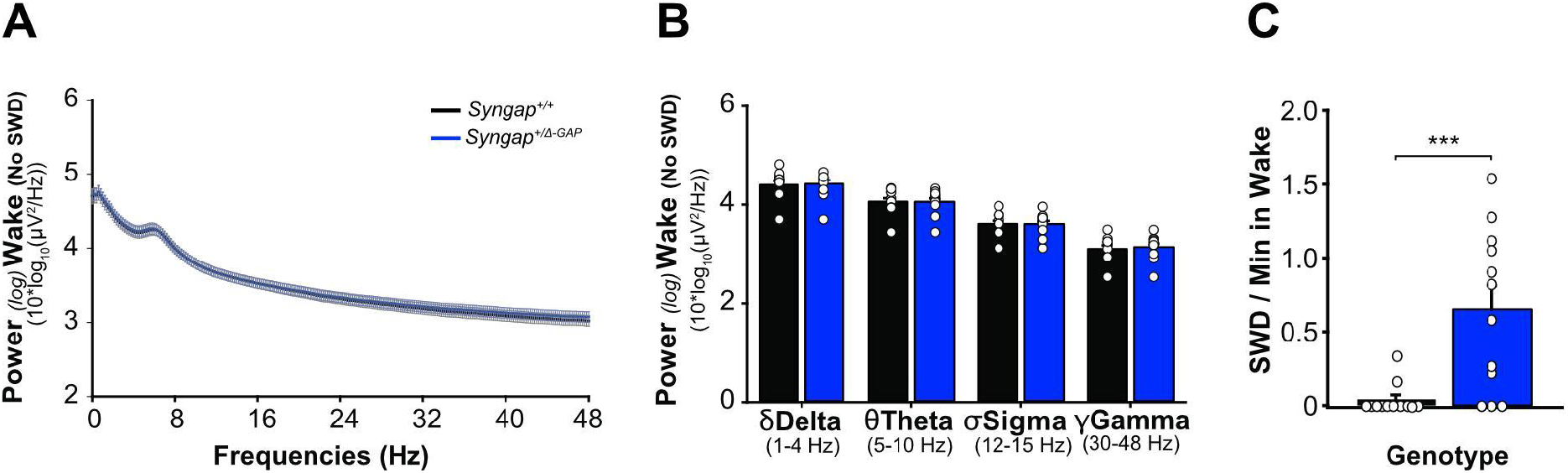
EEG analysis of wakefulness and associated SWD events. Power spectral profile (A) and bands (B) of wake states (excluding SWDs) during wakefulness are comparable between *Syngap^+/Δ-GAP^* and WT littermates. (C) Ratio of SWD events per minute of wakefulness is significantly greater in *Syngap^+/Δ-GAP^* rats compared to WT. (n_+/+_ = 12, *n_+/Δ-GAP_* = 12).

**Supplementary Figure 7.**
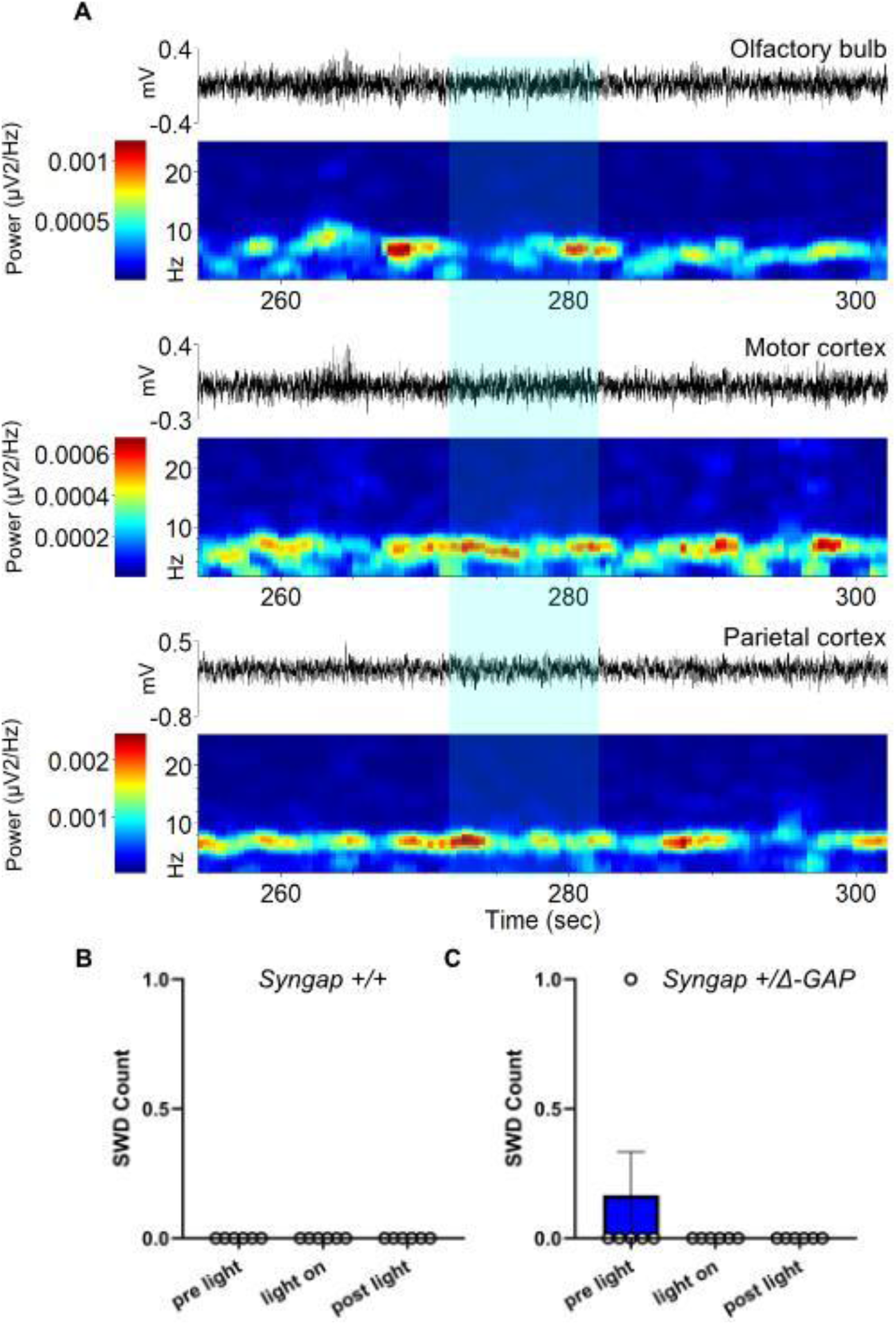
Flashing lights do not drive SWD. (A) Example EEG recording traces and associated spectrograms from a single *Syngap^+/Δ-GAP^* rat before, during and after exposure to flashing light stimuli. (B, C) Average SWD event count per genotype for the 10 sec before (pre), during, and after (post) light exposure (WT left; *Syngap^+/Δ-GAP^* right). Blue bar shading duration when flashing light was on. (n_+/+_ = 6, *n_+/Δ-GAP_* = 6). *mean* ± SE is noted. For *Syngap^+/Δ-GAP^*: one-way ANOVA F_(2,15)_ =1.00, *p*=0.391).

